# PICASO: Profiling Integrative Communities of Aggregated Single-cell Omics data

**DOI:** 10.1101/2024.08.28.610120

**Authors:** Markus Joppich, Rafael Kramann, Sikander Hayat

## Abstract

Various single-cell modalities covering transcriptomics, epigenetic and spatio-temporal changes in health and disease phenotypes are used in an exploratory way to understand biological systems at single-cell resolution. However, the vast amount of such single-cell data is not systematically linked to existing biomedical data. Networks have previously been used to represent harmonized biomedical data. Integrating various resources of biomedical data in networks has recently received increasing attention. These aggregated networks can provide additional insight into the biology of complex human diseases at cell-type level, however, lack inclusion of single cell expression data. Here, we present the PICASO framework, which incorporates single-cell gene expression data as an additional layer to represent associations between cell types, disease phenotypes, drugs and genes. The PICASO network includes several standardized biomedical databases such as STRING, Uniprot, GeneOntology, Reactome, OmniPath and OpenTargets. Using multiple cell type-specific instances of the framework, each annotated and scored with their respective expression data, comparisons between disease states can be made by computing respective sub-networks and comparing the expression scores between conditions. Ultimately, these group-specific networks will allow the identification of relevant genes, processes and potentially druggable targets, as well as the comparison of different measured groups and thus the identification of group-specific communities and interactions.

## INTRODUCTION

Variousl single-cell omics studies are now available from multiple organs and diseases [1–7]. These studies give unprecedented insights into cellular heterogeneity and dysregulated processes in health and disease by comparing different conditions, e.g. disease severities or regions[8]. While all these studies report important mechanistic properties, identifying the key underlying associations to other biomedical data such as related drugs, and diseases involves a lot of manual effort. Computational tools like differential gene expression, PCA or MOFA[8,9] can help to identify condition-specific genes, but these analyses do not yet include systematic integration with other biomedical data that can be used for data-driven prioritization of gene communities for hypothesis generation. Yet, this advanced analysis of single-cell data is one of the grand challenges in single-cell data science[10,11]. Identifying potentially disease associated genesets and understanding their target biology requires association of genes identified in single-cell data analyzes with other biological datasets to capture the relationship between these genes, and diseases, cell-types, protein interaction networks, and their relevance to disease phenotype.

Biomedical data is available in standardized databases such as OpenTargets[12], STRING[13] and msigdb[14]. These databases contain essential information about gene-disease, gene-drug, gene-pathway, subcellular localisation, gene-transcription factor and protein-protein interactions. Such resources have been used to create knowledge graphs such as Bioteque[15], Biocypher[16], PrimeKG[17], or NedRex[18], however, they have not yet been directly linked to single-cell omics data. Common tasks performed on these knowledge graphs are the identification of relevant disease communities for annotated disease, or for drug targets. While some of these frameworks, like NeDRex, allow the incorporation of gene expression values, their focus is still the knowledge graph structure for identifying disease pathways.

In this manuscript we present PICASO, a computational framework that systematically integrates single-cell data with biomedical databases and can be used to identify network communities that are differentially co-expressed or co-regulated in different disease conditions. Our novel PICASO approach for creating and mining biomedical networks to identify explainable disease-associated gene communities and potential drug targets uses gene-regulatory network modeling on the biomedical network representations. Specifically, the PICASO architecture can be used to embed single-cell transcriptomics data within a plentitude of available biomedical databases such as OpenTargets[12], Omnipath[19] (and mirBase[20]), GeneOntology[21], KEGG[22], STRING[13], Reactome[23] and Uniprot[24], and extract condition specific communities and associations. Overall, the full PICASO network consists of 111032 nodes and 1617389 edges collected from the above 7 disparate resources. This embedding primarily scores how well interactions in the knowledge graph are represented by the underlying data. Such derived scores are then used to compare different disease conditions with the help of network topology based approaches available in our PICASO framework. It is possible to identify co-regulated communities and find differential networks (DNETs) between disease conditions.The communities identified by our PICASO method already describe interconnected genes, gene sets, diseases and drugs, but their potential biological relevance, given a disease context, can additionally be predicted from within the framework via large language models (LLMs).

We apply PICASO to the human kidney single-cell data available from the kidney precision medicine project (KPMP)[6] and a multi-modal spatial dataset of human myocardial infarction (MI)[2] to show that PICASO can be used for optimally capturing cell-type and disease-specific gene expression associations across data sources, which can be mined to identify orthogonal molecular, drug and phenotype associations in a cell-type specific manner. In particular, in the MI data, our results reveal multiple communities in fibroblasts and cardiomyocytes, respectively. In KPMP data, PICASO identified various associated communities in all considered diseases, like diabetic kidney disease (DKD) and acute kidney injury (AKI), respectively. Given the large volume of data being generated in the biomedical domain, we anticipate that multimodal computational tools that can capture orthogonal information from different resources will be useful for precision medicine. These can then be used to score the PICASO network to greater detail.

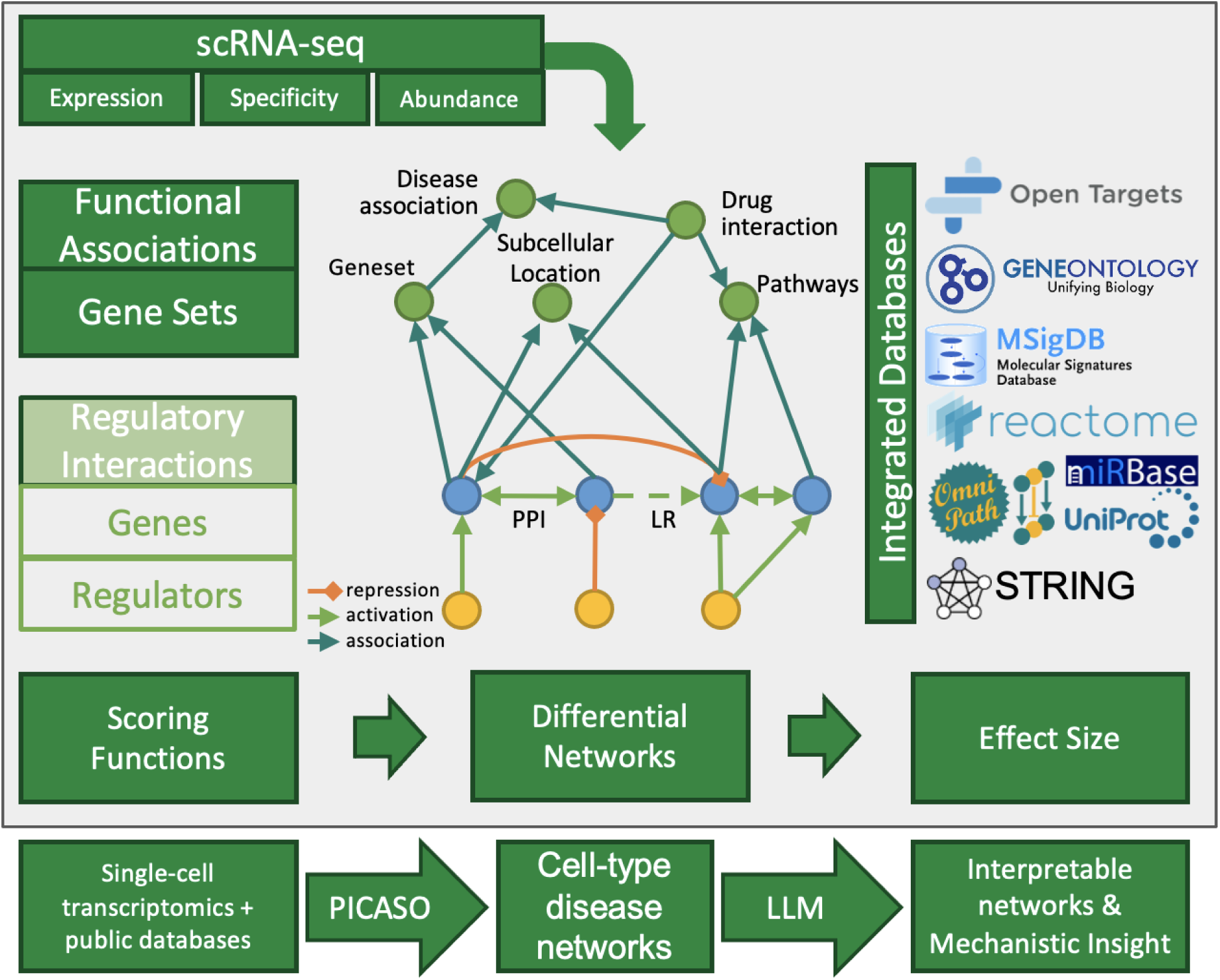

Overview of the PICASO framework. PICASO combines single-cell data with publicly available resources to create annotated networks that can be compared across conditions to identify disease-associated molecular features. An LLM canthen be used to provide automated interpretation, and these features can be used for downstream analyses, further validation

## MATERIALS AND METHODS

### Biomedical data integration and knowledge graph construction

The biomedical integrative network underlying the PICASO framework is a directed graph created by integrating data from various publicly available resources (Figure 1). PICASO is implemented using NetworkX[25]. To create the PICASO network, firstly, data from STRING, Gene Ontology, MSIGDB, Reactome, OpenTargets, human TF and Omnipath is downloaded and aggregated in a unified network (Table 1). This data is represented as a graph, with associations (e.g. interaction, regulation) represented as edges and the different biomedical entities (e.g. genes, diseases) represented as nodes. The following node types are included in the PICASO - genes/proteins, transcription factors, non-coding RNAs (ncRNA), micro-RNAs (miRNA), long-non-coding RNA (lncRNA), pathways, processes, diseases and drugs. These entity types are kept as node type “*property”*. Edges between nodes can currently have only one edge type, such as *interacts*, *part_of*, *relevant_in*, *activates*, *affected_by*, *represses* and *target_of*. The edge type is assigned as indicated by the original database, or simplified to match above-mentioned types. Protein-protein interactions are added to the ‘PICASO network using the STRING database[13]. In the default setting only evidence annotated as fusion, coexpression, experiments, database and text mining are included. Additionally, only interactions with a high confidence (score > 0.7) are kept. All added interactions are assigned “interact” as their edge type category. The PICASO network also contains transcription-factor-target-gene interactions from OmniPath [26]. Here, edges are added with type *activates* or *represses*, depending on the annotated interaction. Transcription factor nodes are annotated as such using the TF node type. Moreover, GeneOntology[21] is added to the PICASO network to include standardized associations covering cellular components, molecular function and biological processes. Edges between GO terms are added as type *part_of*, and edges between genes and their corresponding GO terms are added according to the annotation interaction type by mapping them onto the edge types. Pathway resources such as Reactome[23] and MSigDB[14,27] are added by assigning Gene-pathway edges as type *part_of*. For MSigDB, currently only *Hallmark* and *Curated* gene sets are added. The OmniPath database[28] is also incorporated into the PICASO network. Here, edge types are assigned based on the reported function classified as stimulating or inhibiting. As OmniPath contains different kinds of entity IDs, depending on the reported interaction *type,* genes are added as ncRNA and lncRNA (for lncrna_post_transcriptional) or as miRNAs if the IDs match a known miRBase ID[20].

**Figure 1.**
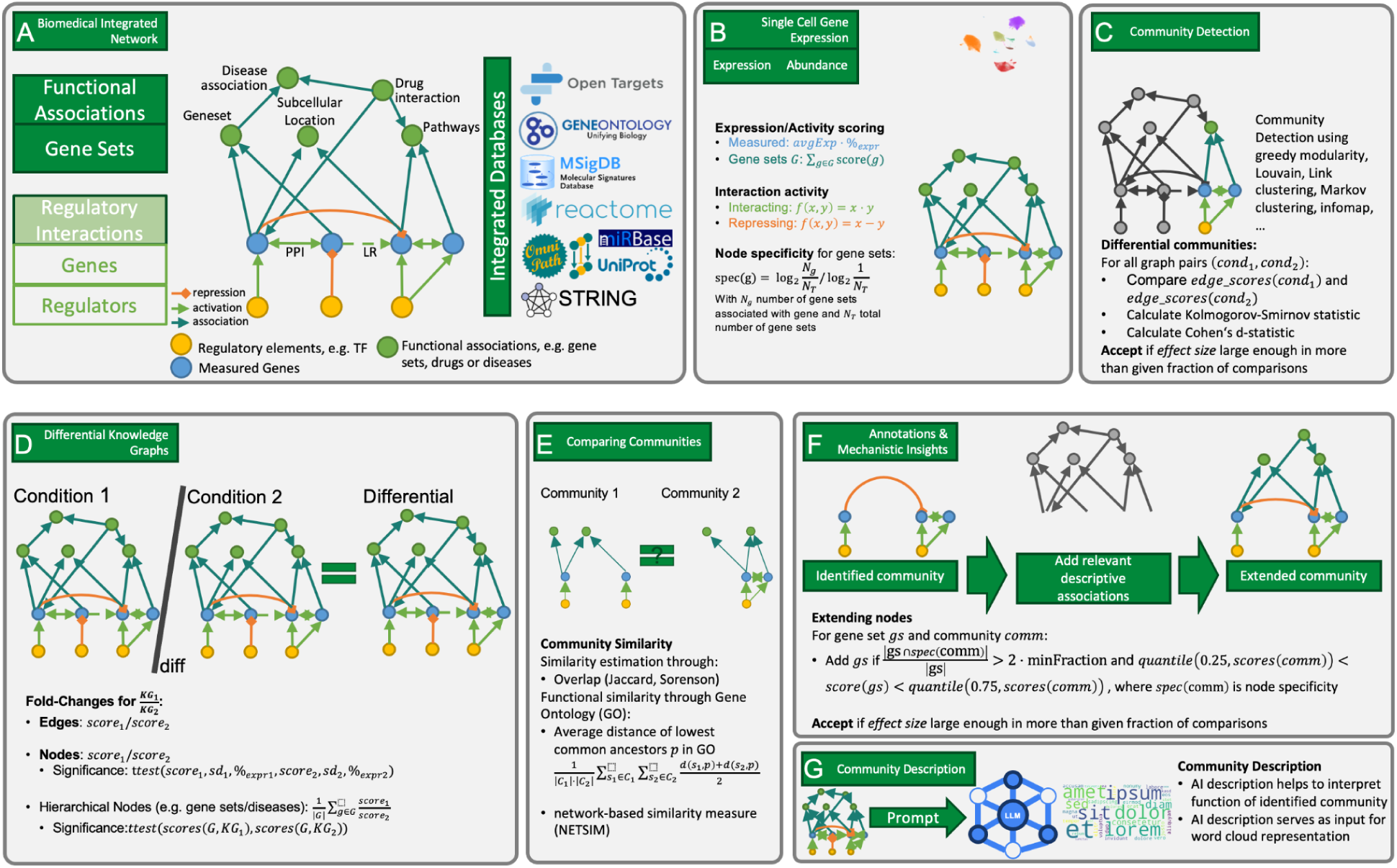
Overview of the PICASO framework. (A) The PICASO network is created from regulatory interactions connecting genes and regulators (e.g. transcription factors or miRNAs), which are further enhanced using gene sets and other functional associations (e.g. disease or drug interactions). The framework integrates several reputable databases for this task (n=8). (B) Given the general PICASO network, gene expression data can be integrated into the network and further propagated along different nodes (e.g. gene sets) and interactions, depending on the kind of interaction. (C) In such an enriched PICASO network several community detection algorithms can be used to identify communities of similar scores (e.g. co-expression). Given multiple conditions, the such identified communities can be compared using multiple measures (KS-test or Cohen’s d-statistic) to only accept such communities which are specific for a specific condition. (D) Another approach to identify condition-specific communities is to first calculate the differential knowledge graph and perform community search within this instance. This effectively searches for co-regulated communities. (E) Particularly in the case of disease conditions or multiple cell types, it is also interesting to see whether there are similar communities across cell types. Therefore the PICASO framework provides functions to compare identified communities by simple overlap or functional similarity. (F) In order to contextualize the identified communities the PICASO framework allows extending the communities to add further relevant descriptive associations (e.g. gene sets, regulators) from the PICASO network. (G) Identified communities can be described using a large-language model (LLM), from which word clouds describing the community’s function are derived.

**Table 1:**
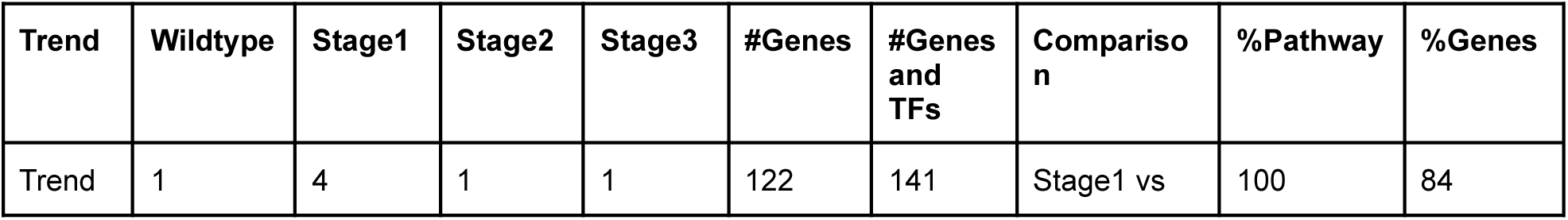
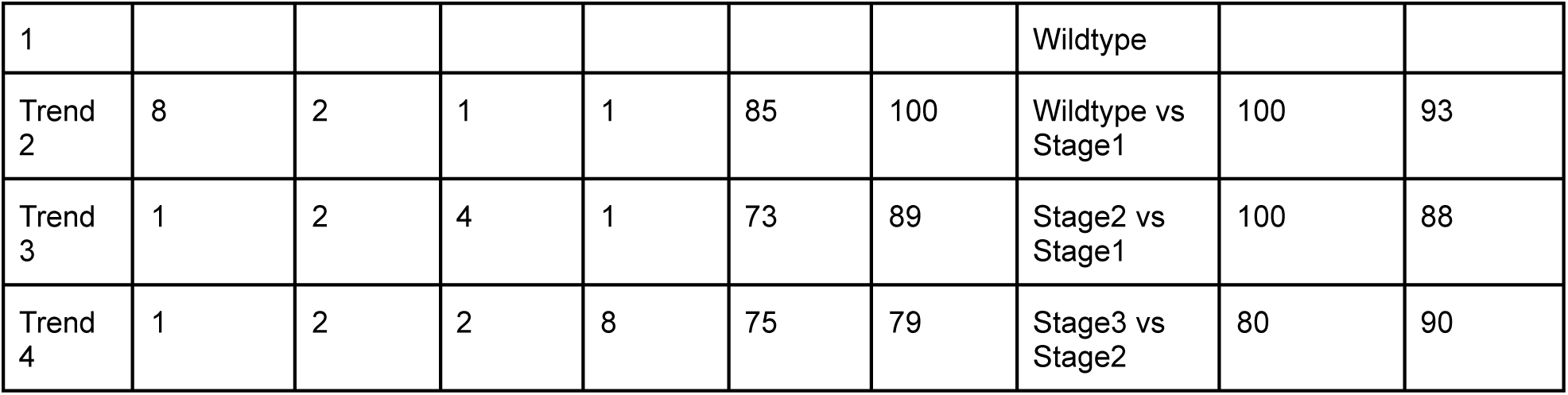
Overview of the simulated trends over the four simulated states (Wildtype, Stage1-3). Trend 2, for instance, has a very high multiplier of 8 in the wildtype state and then gradually declines until no changes of genes associated to this trend will be simulated in stage2 and stage3. While 10 pathways were altered per trend, the number of altered genes ranges from 73 (Trend 3) to 122 (Trend 1). Considering also transcription factors, which induce the changes in the altered pathways, a total number of 79 (Trend 4) to 141 (Trend 1) genes and TFs belong to each trend. The column “Comparison” contains the comparison performed for constructing the differential network. The columns %Pathway and %Genes summarizes the retrieved percentage of pathways and genes, respectively, retrieved by the PICASO method.

Neither stimulating nor repressing miRNA-gene interactions are interpreted as “miRNA represses gene” interaction[29,30]. Finally, disease associations are added from OpenTargets[12]. For each gene-disease association, OpenTargets assigns a *datatypeHarmonicScore*. This score must be 0.8 or higher to be included in the PICASO model. Gene-disease interactions are added as edge type *interacts*. Disease-drug interactions are added as *affected_by* type, and Gene-drug interactions as *target_of* type, respectively. In summary, the PICASO model consists of 111,032 nodes representing genes, genesets, diseases, drugs or cellular sublocalizations, and 1,617,389 edges representing associations between those nodes. Overall, PICASO has 8 node types and 7 edge types collected from 8 resources.

### Single-cell data processing

PICASO assumes that single-cell data has been pre-processed and quality controlled based on standard protocols. Using a QC-ed and annotated single-cell dataset, for all measured features within an Anndata [31,32] object, and for each group, gene expression values are aggregated. For each gene within each group the 0, 25, 50, 75 and 100 percentiles are calculated along with the mean expression. Additionally, within each group, for each feature, the amount and percentage of expressing cells, the group size and the features’ standard deviation is also calculated. These expression-based values are calculated on only those cells which express the gene within the respective group. The resulting table is then written to disk as a tab-separated file as input for inclusion in PICASO.

### The PICASO framework

The PICASO framework provides several functions to access and work with the underlying PICASO network. Statistics on node and edge types can be accessed and visualized using the functions provided. It is also possible to populate the PICASO network with new gene expression data, which were prepared as described above. Nodes for which gene expression data are added also get assigned an additional node type “*measured*”. Using the *GenesetAnnotator* function, it is possible to calculate gene set specificity scores (raw and z-score normalized) for all genes per geneset type (geneset, disease, drug) [33].

The PICASO network is tissue-type and disease agnostic, it only consists of prior knowledge relations from various integrated resources. In order to use this representation for downstream analysis and finding relevant or condition-describing sub-networks, (gene) expression data must be added to the PICASO network. These can then be used to, for instance, describe the activity of nodes or edges in a given condition. Such a scoring of nodes and edges within the PICASO framework is essential for downstream analyses, such as for community identification and constructing the differential PICASO network (DNET). PICASO provides an implementation for calculating node and edge scores within the network by the *MeanNetworkScorer*. Measured entities (such as genes) are scored based on their expression properties. The expression score is calculated as the product of the mean expression with the fraction of expressing cells, summarizing both relevant factors of scRNA-seq gene expression. Subsequently, gene-gene interactions are scored according to a predefined set of scoring functions, which are specific for the respective edge types. The scoring function for *activating* or *interacting* edges is the product of scores, for *repressing* edges the respective scores are subtracted. Gene sets, disease or ncRNAs are scored such that the mean over all their respective gene children is calculated. For any of such nodes, where no genes are found (recursion depth <= 10), these nodes are scored by taking the mean score of all incoming edges. Edges are scored according to their type using the same scoring functions as the gene-gene interaction scoring. Finally, all edge scores are z-score normalized per edge type to account for different edge-type specific characteristics. The calculated scores are used to derive communities from the PICASO network. Additionally, the user can add custom attributes to nodes and edges, e.g. by calculating user-defined scores. Whenever such scores are used in downstream methods, the user can define the attribute-name to be used for collecting edge or node scores. An identified community can be plotted from within the *KGraph* object. Using the *CommunityTool* class within the PICASO framework the user can plot a given community within a set of user-specified PICASO networks, such as different conditions.

### Differential graphs to compare different conditions

To compare different conditions (e.g. disease vs control), there are two methods to identify interesting communities within the PICASO framework. The first one involves the identification of communities in each condition-specific PICASO network. Each identified community is then subset in all condition-wise networks and for each subset the edge scores are collected. A community is then termed differential, if the absolute effect size of the edge scores, between the network in which the community was found and a user-defined fraction of all other networks, is larger than a user-defined minimal effect size. Instead of the effect size the user can also use the Kolmogorov-Smirnov-test statistics from scipy[34] for evaluating differences in edge scores.

The second method to identify condition-specific communities is to first construct a differential PICASO network (DNET). Briefly, given two PICASO networks e.g. obtained from two different conditions or cell-types, for each node and each edge in the two graphs, the log2 fold change between each element’s score is calculated. In order to avoid extreme values, a pseudocount of 0.01 (by default) is used when calculating the log2 fold change. In the default PICASO setup, for measured genes, there is not only an expression score available (e.g. cluster mean expression), but also the number of expressing cells and the standard deviation of gene expression. Thus, for measured genes, in addition to the log2 fold change, a significance value is calculated using the scipy[34] t-test for independent samples, with the node score as mean, the given standard deviation and the number of expressing cells as number of observations. Additionally, the scores for certain node types, like genesets or diseases, are re-evaluated by accounting for the fold-changes of constituent gene nodes. For each of these nodes, first the expression scores are retrieved for all gene children of a node (in both PICASO networks). For each of these genes the fold change between the two PICASO networks is calculated, from which the mean log2 fold change is calculated. Given that matching entities are compared (the same genes within the gene set among both networks), the scipy t-test[34] for related samples is conducted to estimate the significance of the observed change. As input for this t-test the retrieved gene expression scores from both networks and of the genes associated with the geneset are used.

### Community detection

A community in the sense of the PICASO framework refers to a commonly regulated connected set of nodes. Connected means that the regulation of two nodes can be described as a relation: node A up-/down-regulates node B, or simply that node A and B are co-expressed. These relations are important, because within the PICASO method these relations are scored with respect to their plausibility (see section the PICASO framework). Commonly regulated means that the nodes are connected by edges with similar scores. In the regular PICASO network similar scores correspond with similar expression, while in the differential network case similar scores correspond with similar regulation. Putting emphasis on the edge scores is favorable, because then common network-based algorithms for community detection can be used. Within the PICASO framework several community detection methods are already implemented, such as connected components or greedy modularity communities, as well as methods like Louvain (for positive and positive+negative edges), Link clustering[35], Markov clustering[36] or infomap[37,38]. The PICASO network being implemented as a networkx graph, any community detection method able to operate on networkx graphs can be used for this purpose.

### Extending community networks for extracting biological insights

While the identified communities define interesting gene communities (e.g. because community detection took place only on the gene nodes), extracting biological insights from these alone is challenging. However, using the genes in the identified gene communities, additional gene sets, diseases or drugs, which are associated with the retrieved genes, can further describe biological interpretability of our detected communities. We thus extend the communities with relevant gene sets or gene-set-like nodes (e.g. diseases or drugs) and thereby add biological insights to the community. There are two approaches to extend a received PICASO network instance (e.g. a selected community) which are implemented in the *NetworkExtender* class. The *extend_network* method infers an extended PICASO network representation considering the induced subgraph with radius 1 (of specified node type) around all nodes of the original network. Each of the such found new nodes is checked whether it is well represented in the original network. Without loss of generality assume that the checked node is a gene set. It is first checked that the gene set is not too large (<100 children, by default) and also not too small (at least 3 children, by default). For small gene sets (< 5 children) at least 40% of the children must be found in the original PICASO instance, and for larger gene sets at least 50% (both values are specifiable). Children are accepted if they have at least a certain (user-defined) specificity for the considered node type. If a node is accepted using these criterions, edges between this node and nodes from the original network instance are added, if these edges exist in the full/reference network instance. These edges must, however, have a high-enough score (default = 1).

The second option can be used to extend specific node types, e.g. drugs or ncRNAs. While these are similar to gene sets, they might exhibit more specific interaction options. Most importantly it is of interest if there exist, for instance, miRNAs, which can regulate genes contained in the PICASO network, particularly if the miRNA is specific for its target gene. Again, all possible connections in the full network are retrieved for all input nodes in the network to be extended. It is then checked that nodes have a score better than a user-supplied minimal score. The score for the edge connecting the new node and an old node must be within the .25 and .75 quantiles of all edges in the graph. If those conditions are fulfilled, it is assumed that the new node is overall similarly regulated like the whole input network and the new edge (and node) is added to the extended PICASO network.

### Comparing communities

The PICASO framework provides 4 ways (Jaccard, Sorensen, community similarity and network-based similarity) to compare communities which have been retrieved using the methods described above. This is of interest because initially the communities are identified specifically within one PICASO network but similar (differential) communities could also be retrieved from other input PICASO networks. First, the overlap between the different communities can be calculated and shown as a dotplot, where the size of a dot represents the fraction of shared nodes.The similarity of communities can also be assessed in terms of the Jaccard or Sorensen scores and be visualized in terms of a network, where nodes represent communities, and edges gene intersections.

Another method to compare communities is to compare these via ontologies, such as Gene Ontology (GO). With the lowest common ancestor (lca) method, first relevant GO terms for each community are retrieved, and all pairwise lca similarities computed. A GO term is relevant for a community if it is among the most common GO terms (user-defined) and is connected to at least one percent of all community genes. The similarity of two GO terms is defined as the inverse of the average distance of both terms to their lca. The similarity of two communities is calculated as the average weighted sum over these pairwise similarities. The weight is defined as the product of the frequency of the Gene Ontology term per community. Again, these similarities are displayed as a network, where nodes represent communities, and edges represent community similarity.

As an extension, the semantic similarity between two communities is defined as the weighted semantic similarity between descriptive Gene Ontology terms per community, as described above. As semantic similarity the network-based similarity measure NETSIM[39] is used, with a customizable confidence score per edge in the PICASO network. By default, this confidence is set to 1 if an edge is present in the underlying PICASO network.

### Two-level Differential Analysis using the PICASO framework

The PICASO framework can be used with both cell type information and disease condition status to perform two-level differential analyses. For the two-level differential analysis it is assumed that a two-level hierarchy (e.g., cell type and disease status) of input PICASO networks is available. The first level describes the first dimension over which analyses are carried out. This is typically cell-type annotation. The second level contains, for instance, different disease conditions which are compared within each of the first levels.

Following this, for each condition (or zone), communities are calculated independently using the louvain method with the default resolution (set at 4) using the gene network-representation of the PICASO network. These communities are then checked whether they are uniquely enriched in one condition with an effect size difference of at least 1.0 units. Extended communities are only returned if at least one geneset or disease could be associated with the identified community (using the *extend_network* function) to obtain a functional annotation association for each identified community. For gene sets, the *min_gene_spec* parameter is set to 0.5, meaning that a gene for the respective gene set must have a geneset specificity of 0.5 or more to be counted. In addition, the gene sets may have at most 200 members (*max_geneset_size*), and at least 60% of the gene set’s nodes must be in the community for large gene sets (> 5 genes, *min_fraction_large*) or 50% for small genes (*min_fraction_small*). Likewise, these parameters are *min_gene_spec* = 0.8, *max_geneset_size* = 100, *min_fraction_large* = 0.7 and *min_fraction_small* = 0.6 for associating disease target genes with the community. Following this, nodes with drug and ncRNA as nodetype are extended using the *extend_nodetypes* function, such that their specificity is at least 1.0 for drugs and 0.7 for ncRNAs. This functionality is bundled in the *TwoLevelDifferentialAnalysis* class which is used to analyze the data.

### Functional description using LLMs

Functional description of a retrieved community(or simply a gene list) is possible using the *AIDescriptor* class. This class takes as input a path to a storage location for the pre-trained large-language models (LLMs) as well as a model name and corresponding model file to retrieve from Huggingface via the huggingface_hub python module[40]. If the model already exists in the storage location, this model is loaded by the *llama_cpp* python module[41]. The context size is set to 2048 tokens and 16 threads are used for the prediction, by default. The *AIDrescriptor* implements a function *query_genelist*, which takes a gene list as input and a context descriptor (e.g. kidney fibroblasts) for use in the following prompt: *Use the following pieces of information to answer the user’s question*.

*If you don’t know the answer, just say that you don’t know, don’t try to make up an answer*.

*Question: The following genes are dysregulated <CONTEXT>: <GENES>. How are these genes connected and which molecular functions are altered?*

*Do not repeat functions of single genes*.

*Only return the helpful answer. Answer must be concise, detailed and well explained. Helpful answer:*

Where <CONTEXT> is replaced with the context description and <GENES> by the gene list. The resulting answer from the LLM is then returned to the user. The user can also choose to generate a word cloud (using the python package Wordcloud[42]) from the answer, in which the frequency of a word defines the size of the word in the word cloud. For this purpose it is relied on ignored words from the package itself, but also on a custom list of words which occur frequently in the answer without giving any insight on the context of the gene lists, like “genes, important, major, function, cell, protein” etc.

### Benchmarking the PICASO method

The PICASO method is benchmarked on simulated scRNA-seq data. In order to generate a data set with known perturbations, a scRNA-seq data set was simulated using SPARSim[43]. Here, we chose SPARSim as it allows to alter gene expression for certain genes and simulates not only a higher average expression, but also higher frequency of expression within all cells. For each gene within a reference dataset, SPARSim records an average intensity. Most genes have very low intensity values, and therefore a very low expression. Target genes, which are meant to be altered, are thus selected to have an intensity larger than 0.05. This way, only the 3240 genes with highest intensities are target genes for expression perturbation.

In total, we simulated four different states, a wildtype state, and three trends (trend 1 to trend 3) in which the gene expression is perturbed. For these states, the SPARSim multipliers of selected genes are altered in order to modulate changes in gene expression (Table 1). Since PICASO can be used for the identification of altered gene programs, such altered programs also need to be simulated. Using the MSigDB[14] (Version 2023.2) curated gene sets as a repository for such gene programs, it must now be determined which programs are suitable for alteration. Suitable programs have more than one gene and at least 40% of the program’s genes are among the 3240 high intensity target genes. Then, for each trend ten programs/gene sets are selected from all suitable programs. For this, the unsorted suitable programs are iteratively gone through and assigned to the first trend which still has less than ten programs assigned in a way, such that the program’s genes do not overlap with the other trends. In the end, all trends were assigned 10 programs and between 73 and 122 genes.

For each of the four trends, regulating transcription factors (TFs) are annotated. Using the tftargets^1^ GitHub repository and the therein available TRRUST targets[44], regulating TFs are annotated in a greedy fashion. For each gene within the trend the TF which regulates the most other targets as well is chosen. If for a gene already a regulating TF has been annotated, no additional TF is selected. In total, between 79 and 141 genes and TFs are regulated within each trend. While genes by definition can not occur in more than one trend, regulating TFs could occur in more than one trend. For the simulation, the multiplier is set to the maximal value in the respective state over all trends. The simulated gene counts are read into a Seurat object, log normalized with scaling factor 10,000, and finally the 2,000 most variable features are called. Upon these, first PCA is performed, and subsequently UMAP embeddings are calculated. In order to process this data set for our PICASO framework, it is translated into the anndata format using the sceasy^2^ library.

### Retrieving simulated changes

In order to retrieve simulated changes, first four PICASO networks for each simulated trend were created. For each of these networks, the respective gene expression data were added. Then gene scores are calculated and then all remaining scores such as the geneset or disease specificity score are calculated. Genes are annotated regarding their specificity for diseases, genesets and ncRNAs. In order to find the changed genes, DNETs are used and calculated according to the comparison given in Table 1. In order to retrieve genes and pathways, the top 300 nodes with the highest fold-changes are taken. For this ranking, disease and geneset nodes are filtered out if they have less than 5 target genes or a fold change significance larger than 0.1. Currently, all other types of nodes are filtered out if their fold change significance is larger than 0.9 to reduce the network size. In order to evaluate the retrieved network, the amount of recovered genes and pathways is calculated as a sensitivity measure. In order to summarize the sensitivity by taking the single pathways into account, the weighted sensitivity is calculated, which is the weighted average over all pathway sensitivities, weighted by the amount of genes in a pathway, such that larger pathways also have a larger influence.

### Myocardial Infarction analysis

Single-cell RNA-seq data on Myocardial Infarction was obtained from Kuppe et al.[2]. The single-cell data were grouped by cell type and zone (fibrotic, ischemic, border, remote, and control zones as defined by Kuppe et al.) in order to aggregate gene expression per gene into percentage of expressing cells, median and mean gene expression and corresponding standard deviation of expressing cells. Finally 55 (c=11 cell types, d=5 zones) instances of the PICASO networks are created for each cell type-zone-combination. Corresponding gene expression data are added to each PICASO network instance, nodes and edges are scored with the *MeanNetworkScorer* and diseases, genesets, ncRNA and drugs are annotated based on the respective gene set specificity of their target.

To derive celltype-zone-specific communities, first the PICASO networks are organized such that for each cell type all zones are available. Then, the reference zone (control) is used to calculate differential networks for all remaining zones of the cell type against the reference zone. With this experimental setup, the Two-Level Differential Analysis is conducted with default parameters. Relevant communities are then further filtered to include at least one annotated drug and sorted by mean edge score. The shown communities are among the top 3 fibroblast communities.

### KPMP Dataset preparation and analysis

The snRNA-seq KPMP 1.0 data was retrieved from cellxgene [45]. The single-cell data were grouped by cell type and disease (acute kidney injury - AKI, covid induced acute kidney injury - COV_AKI, diabetic kidney disease - DKD, chronic kidney disease - H_CKD, control reference - Ref) in order to aggregate gene expression per gene per cell into percentage of expressing cells, median and mean gene expression, and compute corresponding standard deviation of expressing cells. Finally 80 (c=16 cell-types, d=5 disease conditions) instances of the PICASO network are created for each cell type-disease-combination. Corresponding gene expression data are added to each PICASO network instance, nodes and edges are scored with the *MeanNetworkScorer* and diseases, genesets, ncRNA and drugs are annotated based on the gene expression specificity of their respective targets.

To derive celltype- and disease-specific communities, first the PICASO networks are obtained for each cell-type and disease condition. Then, the reference condition, here the “Ref” samples, is used to calculate differential networks for all remaining disease conditions of the cell type against the reference. Further downstream analysis is performed as outlined above for the MI use case using the Two Level Differential Analysis workflow.

## RESULTS

### The PICASO biomedical knowledge integration framework: resource integration and operations

Multiple publicly available resources are used to build the *PICASO network*. The underlying base network is created using information taken from STRING_DB for protein-protein interactions and co-expressed proteins. Further resources are included to account for either regulatory interactions or add contextual knowledge to the graph: The Human Transcription Factors database serves as the input database for including possible regulators. Finally, OmniPath adds valuable information about many regulatory and communication patterns. Moreover, functional knowledge is required to interpret the identified regulatory patterns. This is taken from commonly used pathway databases such as Gene Ontology, Reactome, and KEGG. Additionally, OpenTargets is used for drug and disease annotations. Using the UniProt subcellular localization ontology, the contained proteins are annotated by their most likely subcellular location (e.g. membrane bound, exogenous, etc.). Gene symbols are chosen as the common nomenclature to annotate genes and proteins when importing individual data resources along with single-cell datasets. In the current version of the PICASO network, this assumption keeps the graph as simple and explainable as possible ‘genes’ transcripts and proteins are represented by a single gene symbol node (Figure 1A). Edges between nodes can have different type annotations, e.g. standing for repressing or activating interactions. Nodes can be associated to several node types: a gene might also be a transcription factor, or a ncRNA. Using an instance of the PICASO network, the corresponding framework can be used to perform multiple tasks. Given experimental data, e.g. scRNA-seq expression data, these can be used to annotate gene expression on measured nodes and propagate gene expression values to, for instance, gene sets. Likewise, interactions can be scored depending on their interaction type and node specificity can be calculated for genes with respect to specific node types (Figure 1B).

A central part of the PICASO framework is the identification of communities. Using the scored and annotated PICASO network instance, multiple methods are provided to identify communities within the PICASO network. If multiple PICASO network instances are available, e.g. due to multiple conditions, it can be checked whether the identified communities have higher scores within one specific condition, resulting in condition-specific communities (Figure 1C). For the case of multiple, condition-specific PICASO network instances, differential networks (DNETs) are useful, because these operate on (log) fold-changes instead of scores. Finding communities in such a differential KG can be used to identify whole regulatory programs being regulated in one condition (Figure 1D). For the identified communities, the PICASO framework can also compute similarity scores based on simple overlap or even functional similarity (Figure 1E). Identified communities are retrieved based on the underlying scores, and thus, descriptive terms may be disconnected during community detection. Using a network extension approach it is also possible to add relevant descriptive associations, such as pathways, subcellular locations, diseases or drugs, to an identified community (Figure 1F). The identified communities can be described within a user-specified context using large-language models (Figure 1G). This textual description can also be converted into a word cloud.

In order to create an instance of the PICASO network, the user can define which resources should be included into the network. By default, and in all evaluations presented in this manuscript, all available resources are included. This yields a PICASO network instance with 111,713 nodes and 1,569,658 edges (Figure S1 A). Most nodes are genesets (49.7%) followed by genes (37.8%) and diseases (7.0%) (Figure S1 B). Most edges are between genes and gene sets (52%) and between genes (21.3%) (Figure S1 B). It is thus not surprising that most edges are added from KEGG (31%), GeneOntology (22.4%) and STRING DB (18.1%) (Figure S1 C). It is maybe surprising that edges from OpenTargets only make up 5.7% of all edges, given that there are many more edges collected. For the construction of the base PICASO network, however, only gene-disease associations with a score of at least 80% were added. Most nodes have a low number of in-edges, however, particularly gene sets can have many in-edges from their annotated genes. This is, however, given the definition of, for instance, Gene Ontology not surprising. The base PICASO network contains all interactions between genes, genesets, diseases and drugs (Figure 2). At this stage, it can be used to interpret the associations for a given context, e.g. around a specific gene, like the sodium/glucose cotransporter 2 (SGLT2) encoded by the SLC5A2 gene. In the shown 1-neighborhood of the gene, limited to 10 representatives per node type, it can already be seen that SGLT2 plays crucial roles in various diseases, like diabetes, obesity, type 2 diabetes nephropathy, heart failure or non-alcoholic fatty liver disease, against which SGLT2 inhibitors are currently under clinical investigation[46]. Outgoing from the core PICASO network, the PICASO framework offers the possibility to annotate the graph with expression data (e.g. from scRNA-seq data) and to calculate expression scores, which take gene expression levels and expression sparsity into account. These scores can then be used to infer activity scores on other (unmeasured) nodes, such as gene sets, diseases, drugs or ncRNAs. Within the PICASO framework, the score distribution per node type, edge type or subsets of both can be explored. While the expression scores for measured genes can, most likely, be well interpreted - particularly if the input data was properly normalized - this is not the case for the inferred scores or edge scores, e.g. for activating, repressing or interacting edges. Thus, for these edge types, the z-score is calculated on a per-type basis, which makes the different scoring schemes comparable.

**Figure 2.**
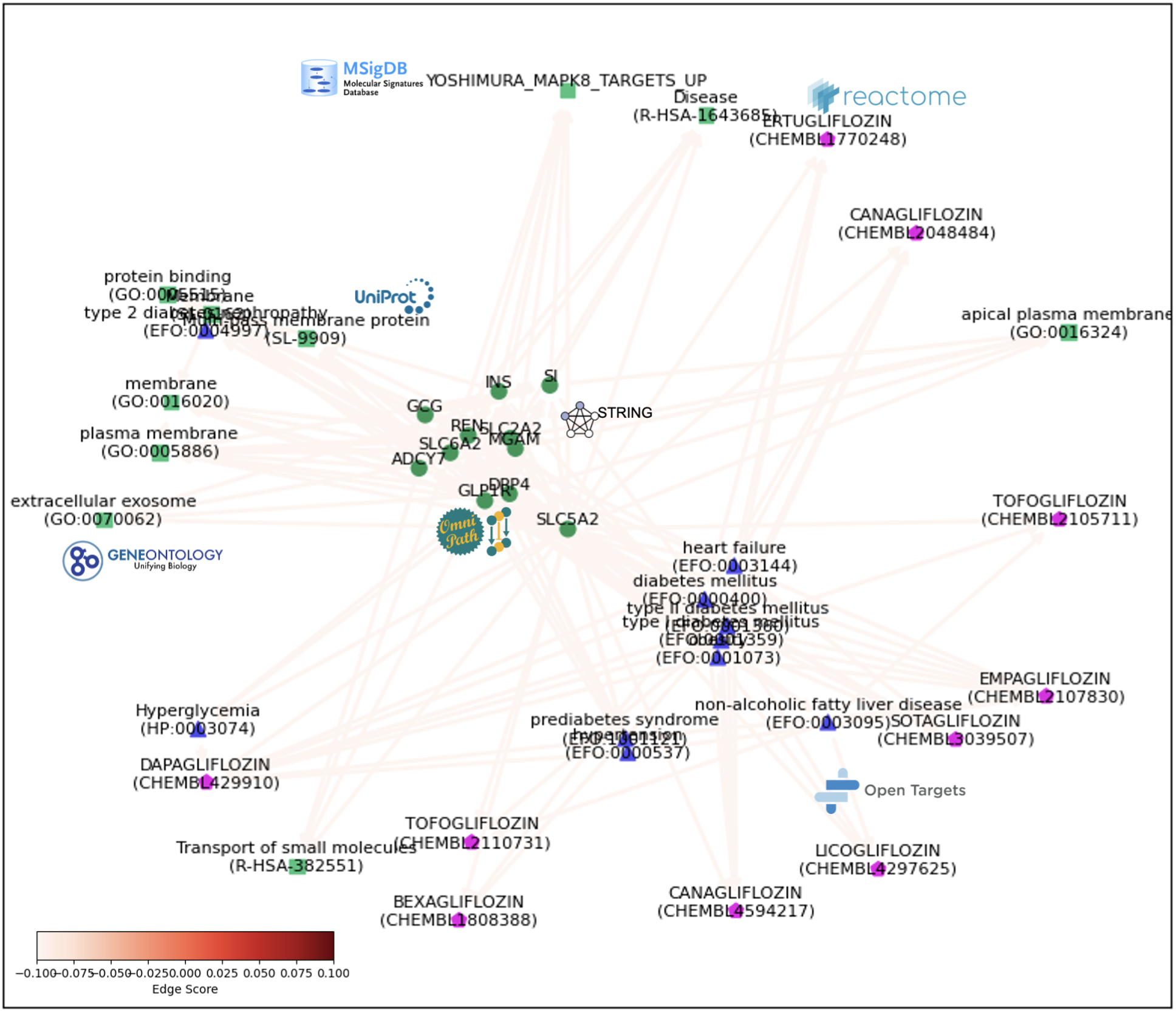
Illustrative example of a PICASO network for the induced subnetwork around the SLC5A2 gene which encodes the sodium/glucose cotransporter 2 (SGLT2) [73]. The cotransporter plays an important role in multiple diseases and is actively investigated as a druggable target[46]. For some nodes, it is exemplarily shown from which resource the respective node and edge are added.

### Benchmarking PICASO using simulated single-cell RNA-Seq data

Given a PICASO network, first we wanted to check whether it is possible to optimally retrieve simulated changes. For this, a scRNA-seq data set was simulated which consists of four distinct groups of cells (Figure 3A). Furthermore, four different gene expression patterns are simulated (for ten randomly selected genesets) over these four groups of cells (Figure 3B). These are described as 4 trends. Using the PICASO framework, the aim here was to see if these changes can be optimally retrieved. After constructing a PICASO instance for each of the four groups of cells, DNETs are constructed as described (see Methods). In order to retrieve the changed genes and pathways for trend1, the differential KG between stage1 and wildtype is calculated. In a first step it was checked whether the resulting log fold-changes can be used to distinguish the several simulated trends. Indeed, the genes associated with trend1 show higher log-fold changes than genes associated with the other trends (Figure S2 A). This is also visible when focusing only on altered pathways (Figure S2 B). Following the calculation of the differential KG, which contains all genes, this representation is filtered to only keep the 300 genes with highest fold changes and removing gene sets which have less than 3 genes associated with. The resulting PICASO instance is then plotted with node sizes corresponding to the node’s log fold change, and edge color to the edge’s log fold change, respectively (Figure S2 C). In terms of being able to retrieve the genes of the altered gene sets and the gene sets themselves, using the described approach, between approx. 80% and 91% of genes can be retrieved, and often all pathways (Figure 3C). The missing genes can be explained as these genes have very low expression and are barely expressed. In case of missing TFs, these can occur in multiple trends and therefore may not exhibit as clear signals as needed for discovery. The missing gene sets do not fulfill the significance threshold for the top 300 nodes. Nonetheless, a high majority of altered genes and gene sets are detected. Most importantly, not only the genes and gene sets themselves are detected, but also the added transcription factors, which are meant to regulate the respective gene sets. This shows that the general approach of encoding gene expression scores in the PICASO network and log fold changes in the differential networks enables the discovery of altered gene expression programs.

**Figure 3.**
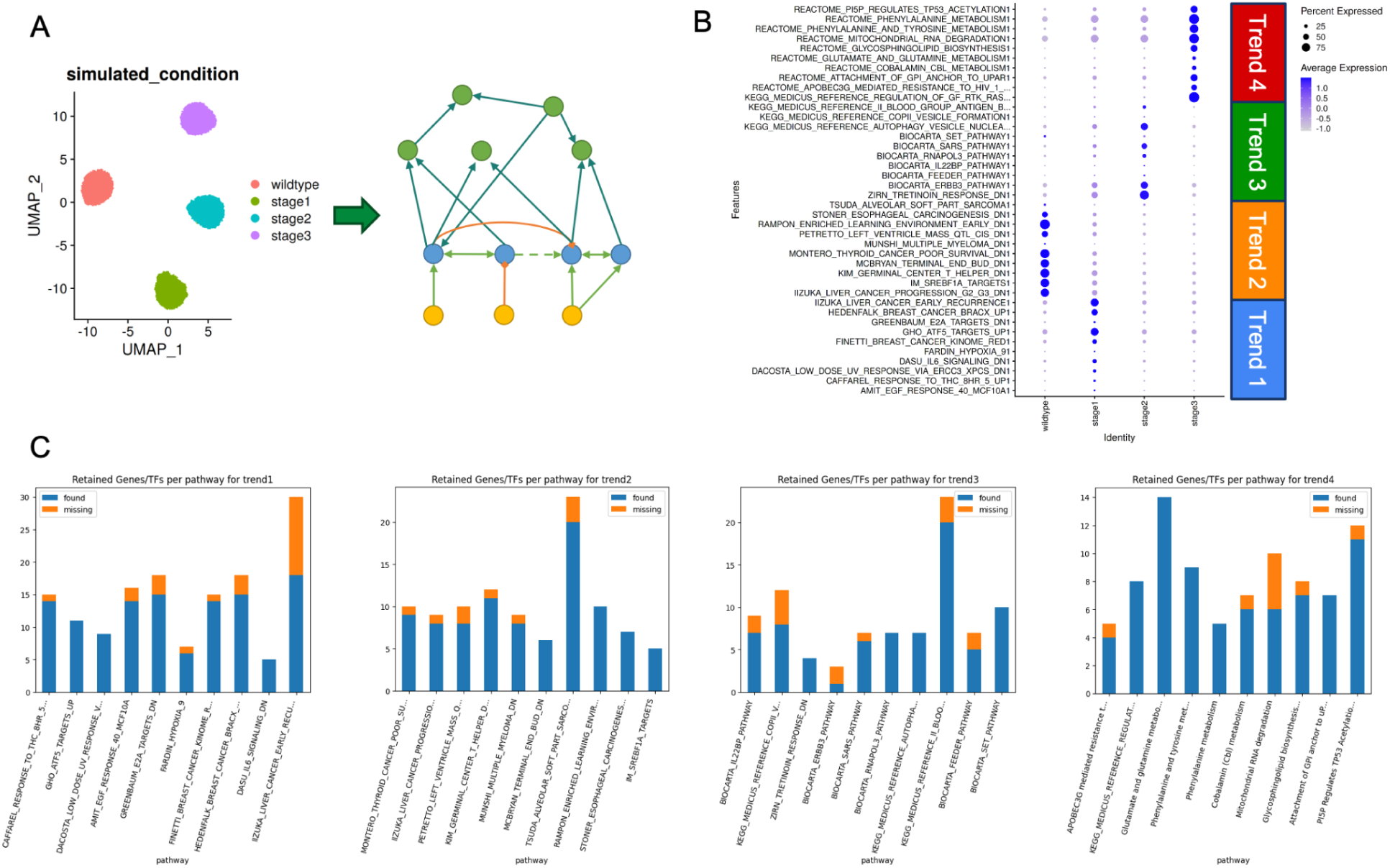
(A) UMAP of the simulated single-cell data. For each stage of the simulated data, a PICASO network is created and network scores have been calculated. (B) The perturbed gene sets show distinct patterns/trends over the different stages. The community scores of the perturbed genesets show distinct expression patterns, e.g. very broad expression in all cells, or even only in few cells. (C) Evaluation of the accuracy of the retrieval of the genes associated with the several simulated trends. While the genes of the simulated gene sets are well discovered, missing entries usually are simulated transcription factors, which are regulating multiple pathways of different trends at once and thus do not show clear signals.

### Disease-associated co-regulation communities identified through PICASO

#### Use case 1: Application of PICASO to human myocardial infarction single-nuclei data

After having shown that the proposed framework detects altered gene regulatory programs (including relevant transcription factors) in the previous section, we apply PICASO to a real-world problem setting using publicly available human myocardial Infarction snRNA-seq data[2]. The dataset consists of samples obtained from five different disease-specific zone of the left ventricle of the human heart, namely - the unaffected control tissue, remote zone (RZ), border zone (BZ), ischemic zone (IZ) and the fibrotic zone (FZ). In total, the data set consists of 191795 cells and 11 cell types. Each cell type was processed separately and transformed into a PICASO network (Figure 4A). Using the two-level differential analysis approach, co-regulated communities are identified. In order to achieve this, a set of differential networks (DNETs) is constructed for each cell type over all conditions, and individually compared each time to the PICASO network from the respective control samples, resulting in several differential communities identified in the different cell types (Figure 4B). As expected, most communities are identified in the ischemic and fibrotic zones. Since the edge scores are used for identifying communities, which correspond to the log2 fold changes between the distinct zones and the control, this overrepresentation of the ischemic and fibrotic zone can also be seen in the distribution of edge scores on those DNETs (Figure 4C). In the heatmap for all identified fibroblast communities (see Methods for parameter settings on identifying the communities), the respective communities are clustered by their score-pattern over all zones (Figure 4D). A total of 28 communities were identified within fibroblasts, 2 of them in the remote zone, 8 in the fibrotic zone and 18 in the ischemic zone. Given that communities are derived in all DNETs independently, analyzing the communities’ similarity in terms of the functional similarity score NETSIM can be helpful to identify functional similar communities over different DNETs (Figure 4E).

**Figure 4:**
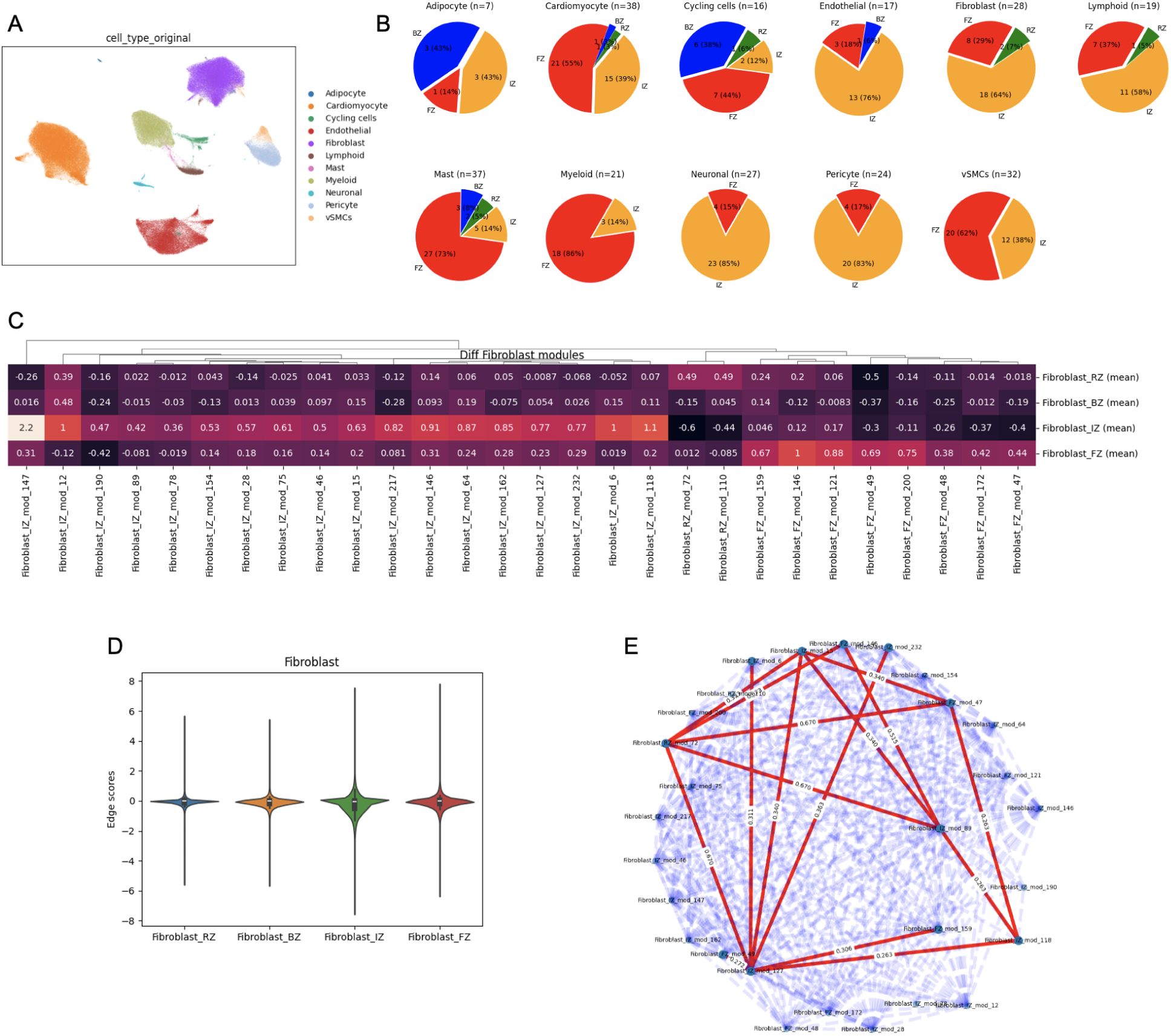
Case-study - Application of PICASO to Myocardial Infarction single-cell transcriptomics data[2]. (A) UMAP of the MI human heart data. PICASO networks for all cell types and all 5 conditions (control, RZ, BZ, IZ, FZ) are created. (B) The resulting count of identified communities in the considered cell types from which the number of communities identified in the comparison of respective zones vs controls can be obtained. (C) The majority of communities are detected in the ischemic (IZ) vs controls, and fibrotic (FZ) zone vs controls comparison. (D) Edge score distributions in the DNETs comparing each zone with the control samples in Fibroblasts. In the IZ and FZ, the distribution of edge scores (log2 fold changes) appears wider than in the other conditions. (E) A network diagram showing the overlap between the retrieved communities in terms of the NETSIM similarity score.

The overview plot for the first community shows the identified and extended nodes colored by node type and scaled by node score (Figure 5A). This community was identified on the gene subset of the FZ/CTRL DNET network. The miRNAs, drugs and GO genesets are annotated afterwards as described for the two level differential analysis workflow (see Methods). The extended geneset associations relate to serotonin clearance and vitamin B6 activation to pyridoxal phosphate.The serotonin clearance is mainly driven by the gene MAO-A, using serotonin as educt, which is already known to be highly expressed in fibroblasts after myocardial infarction[47]. The mechanism leading to MAO-A upregulation is yet unknown, and the derived community might shed light into this process. Furthermore, vitamin B6 activation to pyridocal phosphate is already known to be cardiac-protective[48]. We also identify the drug Disulfiram, which targets the gene ADH1B (alcohol dehydrogenase 1B), which is upregulated in fibroblasts of the fibrotic zone and has been shown to be block fibrosis in human and mouse ocular scarring[49]. Recently, it was shown that disulfiram is linked to cardiac protection through inhibiting gasdermin D[50]. Specifically, inhibition of gasdermin D is linked to reduction in pyroptosis which is a form of cell death involving destruction of cell membranes and the release of pro-inflammatory factors [51].

**Figure 5:**
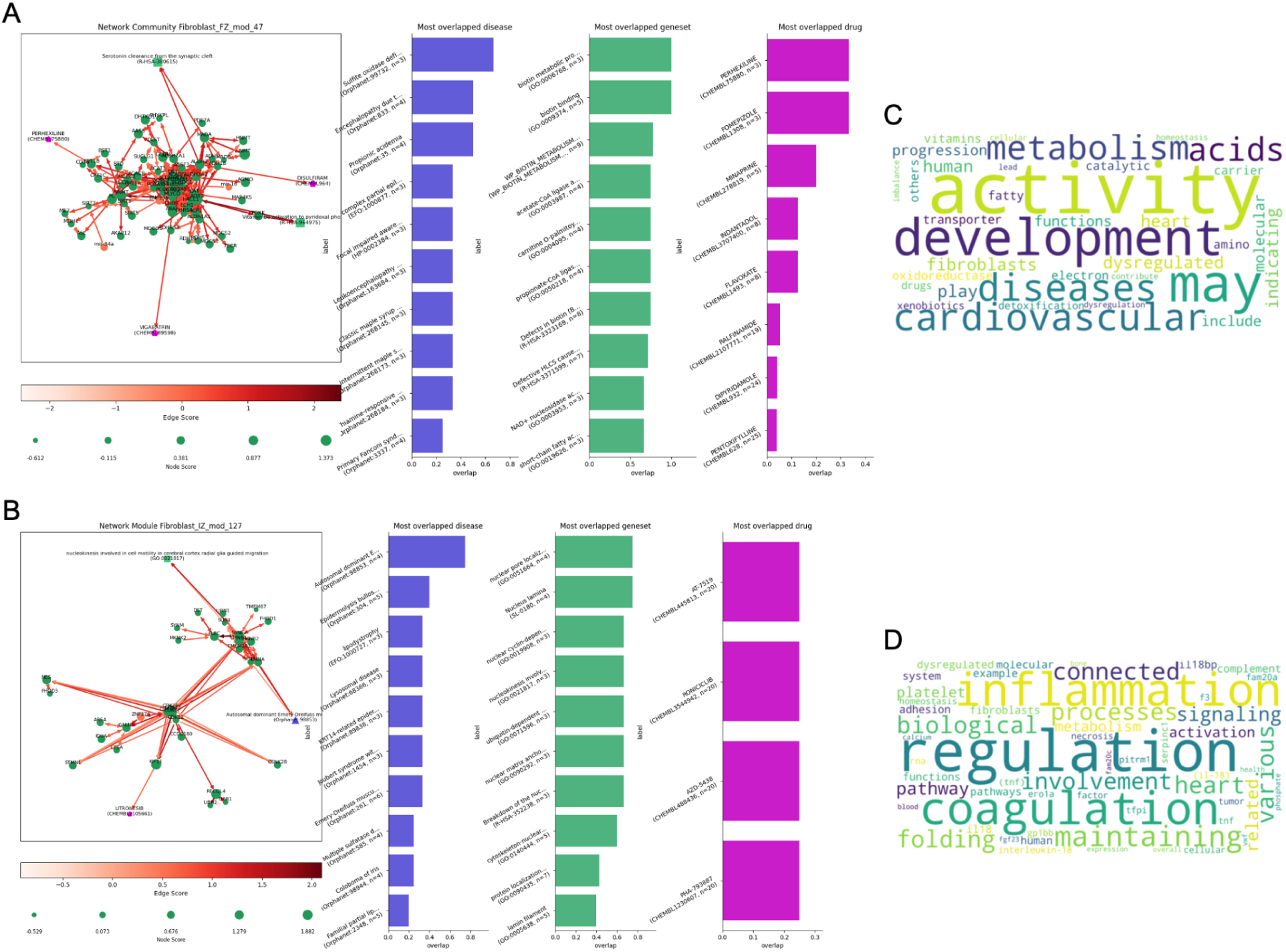
Selected PICASO differential networks identified for Myocardial Infarction. (A) Network overview of the fibroblast community 47 identified in the fibrotic zone vs control comparison and (B) community 127 identified in the ischemic zone vs control samples comparison. The first panel shows the identified community with all nodes sized by score and edges colored by score. The next three panels show the node overlap in the identified community with disease, GO genesets, and drugs. (C) and (D) show the corresponding LLM based AI Descriptor word clouds for the communities shown in (A) and (B), respectively.

Similarly, perhexiline inhibits carnitine palmitoyltransferase I (CPT1A) and has been evaluated to be a viable therapy in patients with refractory myocardial ischemia[52]. The next three panels (Figure 5A) show diseases, GO genesets and drugs most overlapping with those communities. If such a term is not included in our identified network, its target genes were either not specific enough for the term or the overlap was too small (Methods). In general, all identified terms are related to metabolic reactions. Further, the edge scores of the identified community in the samples from the fibrotic zone is higher as compared to others, and the dotplot shows the gene expression quantification in the single-cell data (Figure S3 A). However, it is to be noted that the communities are detected using the edge scores obtained from calculating differential expression in the DNET, and that the dotplot shows absolute fold changes of the nodes (genes). In order to derive more insights into the molecular basis of the community, we provide an *AIDescriptor* class to query a large-language model trained on biomedical literature in order to derive a textual description of the community [53,54]. This textual description is turned into a word cloud (Figure 5C) to make its content more easily understandable. In this use case, the identified community is highly related to metabolic activity.

We illustrate a second identified community in fibroblasts of the ischemic zone (Figure 5B). First it can be noted that the community appears to consist of two subnetworks which are well connected. The one subnetwork contains genes associated with “nucleokinesis involved in cell motility in cerebral cortex radial glia guided migration” (GO:0021817) as well as “Autosomal dominant Emery-Dreifuss muscular dystrophy” (Orphanet:98853). Both diseases are supported by overlapping disease and geneset annotations. The connection between these two entities and cardiovascular diseases has already been shown in aggregated resources like malacards[55]. The identified community consists of CDK12/13/17/16 genes (cyclin-dependent kinases), which on their own were recently identified to be important regulators on cardiac stress[56]. CDKs are known druggable targets, and our community analysis identified litronesib, which is a Mitosis-Specific Kinesin Eg5 inhibitor[57]. Interestingly, roniciclib, one of the overlapping drugs, is also a well known pan-CDK inhibitor[58]. In the cumulative histogram it can be seen that the presented community is enriched in the ischemic zone, which is confirmed by the dotplot of the contained genes in the underlying scRNA-seq data (Figure S3 B). Using the *AIDescriptor*, the employed LLM (Figure 5D) highlights the role of this community to terms closely related to fibrosis such as inflammation[59] and coagulation[60].

#### Use case 2: Application of PICASO to Human Kidney precision medicine project single-nuclei data

Kidney precision medicine project (KPMP) data consists of multi-modal human kidney data from hypertensive chronic kidney disease (H_CKD), acute kidney injury (AKI), covid acute kidney injury (COV_AKI), diabetic kidney disease (DKD) and controls[6]. We used the single-nuclei data including 200,338 cells to build a PICASO network for each cell-type per disease (Figure 6A). Subsequently, the differential networks (DNETs), with the control samples as reference were calculated for each disease and cell type. Most differential and interesting PICASO communities are found within the AKI and COV_AKI conditions, with the exception of PapE cells, where most communities are detected in the DKD samples, and the NEU cells, where most communities were found in H_CKD (Figure 6C). In the remainder of this section we focus on endothelial cells (EC). In the distribution of edge scores, which correspond to the log2 fold changes of the DNET between each disease condition and the reference, a shift towards positive log2 fold changes is observed (Figure 6B). This shift is particularly strong in COV_AKI. Notably, DKD has many down-regulated genes. The prevalence of COV_AKI associated communities is shown in the heatmap comparing all identified endothelial cell communities (Figure 6D). For a more detailed analysis we focus on the only H_CKD associated community (EC_H_CKD_mod_14) as well as one COV-AKI associated community (EC_COV_AKI_mod_29), which was selected to highlight the functionality of PICASO to discover drug overlap results.

**Figure 6:**
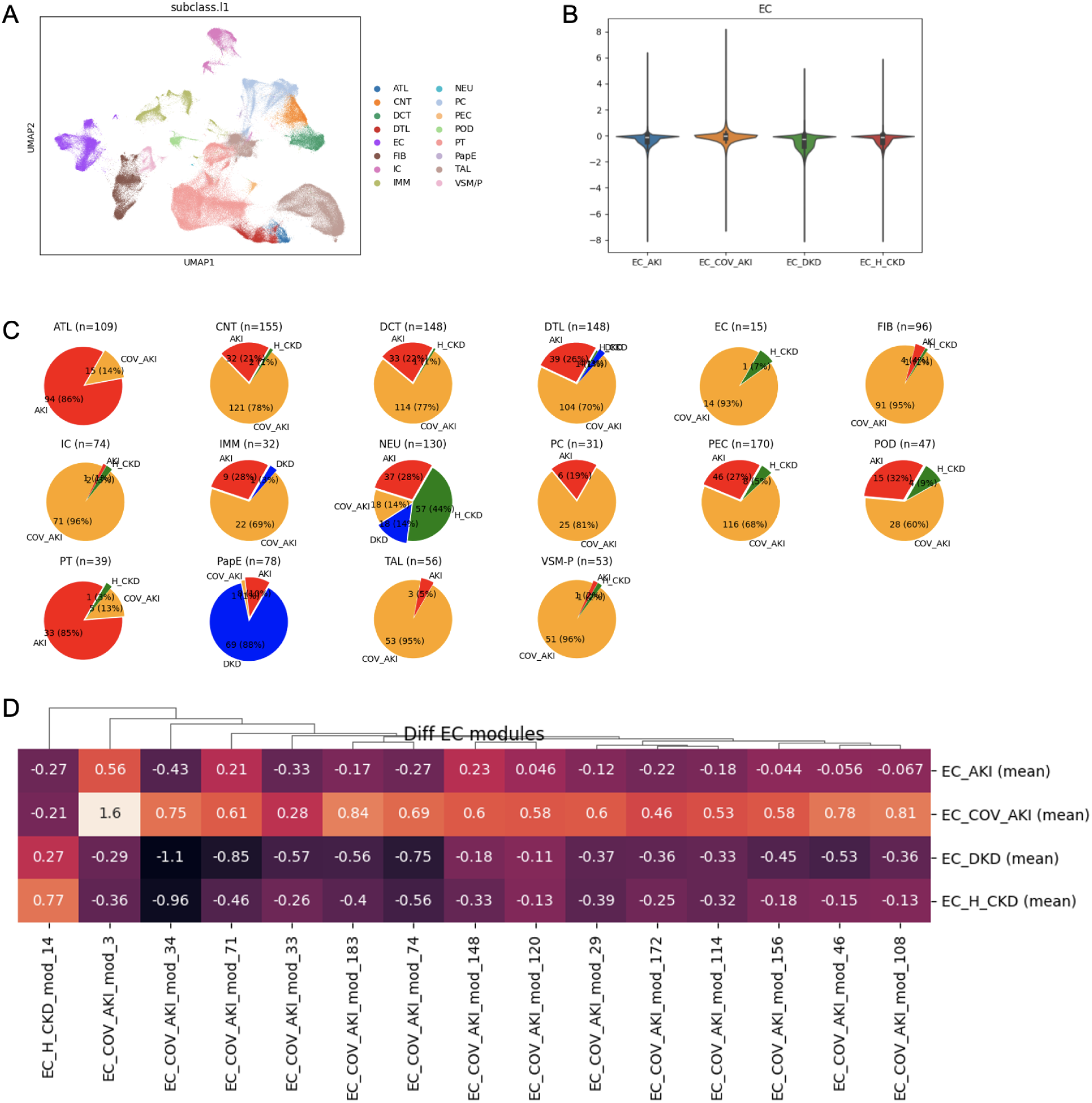
Case-study kidney diseases in endothelial cells using KPMP single-cell transcriptomics data. In the co-regulation use-case for the kidney precision medicine project data (A) PICASO networks for all cell types and all 5 disease conditions (reference, AKI, COV_AKI, DKD and H_CKD) have been created. (B) The resulting number of identified communities in the considered cell types are shown as pie charts, from which the number of communities identified in the respective zones can be taken.The majority of communities is detected in AKI and COV_AKI. (C) Edge score distributions in the DNETs comparing each zone with the control samples in endothelial cells. In COV_AKI the distribution of edge scores, which here are log2 fold changes, appears shifted towards positive log2 fold changes, in contrast to DKD, where a shift to negative log2 fold changes can be seen. (D) Heatmap showing the endothelial cell associated differential communities with their mean edge score per disease. As seen in the pie charts, most communities are found in COV_AKI.

EC_H_CKD_mod_14 (Figure 7A) is associated with the Bartter syndrome (Orphanet:112) and Potassium transport channels (R-HSA-1296067). Both observations are, however, linked: Bartter syndrome is a disease of renal electrolyte transport, often already observed in neonatal patients[61]. Mutations in Na-K-2Cl co-transporter encoded by SLC12A1 are associated with Bartter syndrome[62]. Furthermore, mutations in the renal potassium channel ROMK encoded by gene KCNJ1[63], renal chloride channel C1C-Kb encoded by gene CLCNKB[64] and Na-Cl co-transporter encoded by gene SLC12A3[61] are also associated with this disease. All of these, but SLC12A3, are contained in the network identified by PICASO. The potassium channel ROMK is also annotated as such via the reactome geneset R-HSA-1296067. In the overlap analysis regarding diseases and genesets the identified Bartter syndrome and potassium transport channels are confirmed.

**Figure 7:**
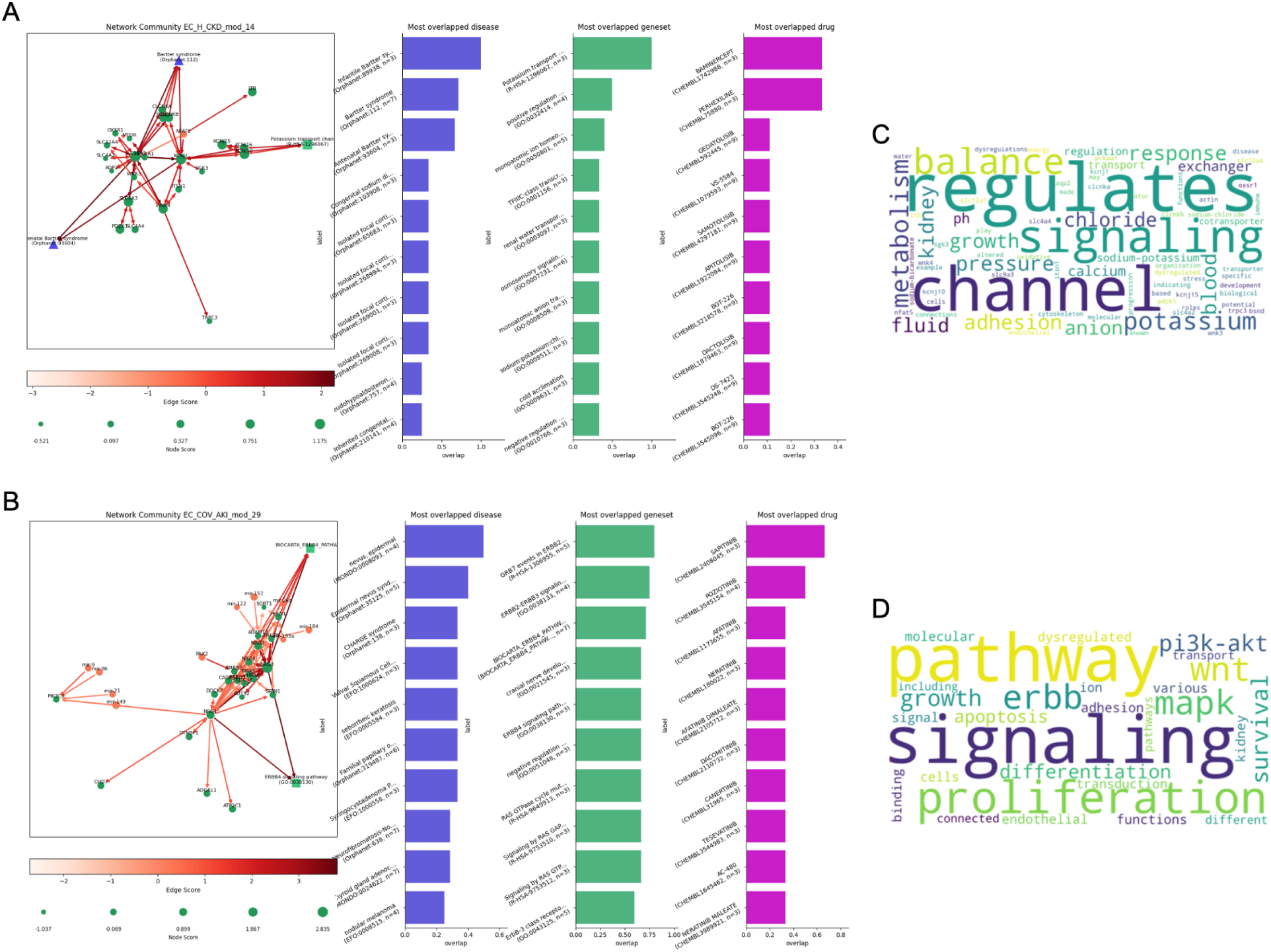
PICASO selected differential networks in H_CKD and COV_AKI kidney conditions in endothelial cells using KPMP single-nuclei transcriptomics data. (A) Network overview of the Endothelial cells (EC) community 14 identified in H_CKD vs control comparison and (B) community 29 identified in the COV_AKI vs control samples comparison. The first panel shows the identified community with all nodes sized by score and edges colored by score. The next three panels show the node overlap in the identified community with disease, GO genesets, and drugs. (C) and (D) show the corresponding LLM based AI Descriptor word clouds for the communities shown in (A) and (B), respectively.

While there is no drug directly annotated in the identified community, the overlap analysis reveals two associated drugs (one gene of three overlapping), namely baminercept and perhexiline. Both only share one of three associated genes in the network. Baminercept[65] inhibits Lymphotoxin-β, which is encoded by the LTB gene and allows phagocytic cells to bind to endothelial cells[66]. We also observe that compared to the reference condition, the expression of LTB is increased in H_CKD. In general, the identified community is only slightly enriched in H_CKD compared to the other disease conditions, which can be seen from the cumulative histogram on the edge scores as well as the gene expression data in the dot plot (Figure S4 A). The AIDescriptor captures well the relation of the community to the ion channels and through these with metabolism (Figure 7C).

Finally the COV_AKI associated community EC_COV_AKI_mod_29 (Figure 7B) is interesting because it is associated with the ERBB4 signaling pathway, which is well supported also by the geneset overlapping analysis and existing literature[67]. No disease or drug is extended in the network community, but nevus syndromes are associated through the HRAS and NRAS genes in the community. These are, however, not up-regulated in the differential expression analysis. ERBB4 signaling in endothelial cells has been previously associated with cardiac fibrosis following myocardial infarction[68], hence it could be that a similar function is also found in the kidney[67]. Being able to interfere with this ERBB4 mediated cellular crosstalk might be an option to avoid the detrimental effects. Similarly, it has also been shown that ERBB4 plays an integral role during SARS-CoV-2 infection, during which sapitinib, the most overlapped drug, suppresses SARS-CoV-2 infection, while blocking ERBB4 signaling as a pan-ERBB inhibitor[69]. From the cumulative histogram of the community it can be seen that only half of the edges diverge from the edge scores in the other conditions (Figure S4 B). By looking at the dotplot of the gene expression in the underlying snRNA-seq data it can be hypothesized that this change in edge scores is mainly driven by expression of NRG3. Given that NRG3 is the ligand for ERBB4, being able to control the gene expression of NRG3 might be an interesting target for inhibiting the auto-activation of the ERBB4 pathway. The *AIDescriptor* generated word cloud on this community correctly describes the community as being about the ERBB signaling pathway, leading to proliferative behavior of the endothelial cells, if not counteracted using pan-ERBB inhibitors (Figure 7D).

## DISCUSSION

A rapid development of single cell sequencing technologies has been seen in the last few years. In the beginning the pure description of novel single cell dataset was of high interest, nowadays it is more commonly tried to interpret such data sets mechanistically. Recently a lot of attention was paid to describing new tools which infer gene regulatory networks from single cell RNA-seq data [70,71]. Despite the many existing databases on gene regulatory elements or even gene/protein-gene/protein interactions, this information is not used by most of these methods. From the integrative bioinformatics perspective not using such information is a major disadvantage of these methods.

The PICASO framework describes a novel approach to represent (single cell) gene expression data in a network built from prior knowledge and provides new approaches to interpret this data. By leveraging gene expression data into the PICASO network structure it is possible to apply graph-based methods to infer novel insights into the data. A current limitation of PICASO is that it relies on prior knowledge edges and contextualizes these by checking or scoring how well these edges are represented in the underlying data. For the future, combining PICASO with methods for deriving gene-regulatory networks appears to be a logical extension. Thus, any insights through PICASO do not result from identifying possible new edges, like in traditional knowledge graph tasks, but result from checking or scoring how well an existing edge is represented in the underlying data. Such an approach of going from simple gene sets to graph-based interpretations is not completely new. In GGEA[72], for instance, the authors extend existing gene sets with a generalized gene regulatory network and use these augmented “graphs’’ for further analysis. With our approach we do not rely only on one kind of gene regulatory network, but incorporate several interaction networks, be it of gene regulatory nature or protein interactions. Moreover, our analysis does not try to explain existing gene sets only, but finds communities in a data-driven way. This way it is possible to find co-expression communities, or co-regulated communities from the novel differential network (DNET) concept, which may consist of multiple gene sets at once and thus represent larger connections than a single gene set alone. Further existing knowledge represented in the PICASO network, such as disease or drug annotations serve then to contextualize the identified communities for best interpretability.

With the PICASO network and methods it is possible to represent expression data within existing knowledge, while combining this with data-driven analyses. This enables the user to generate hypotheses in a data-driven manner, which can be further made understandable by network extension approaches, gene set overlaps or AI-powered descriptions of output nodes using large-language models.

Using simulated data, we show that our framework is able to identify perturbations within scRNA-seq data, of which a significant amount of changes could be traced down to the single gene. While not all perturbed genes could be identified, likely due to too small or non-consistent regulations, e.g. if a transcription factor is relevant for multiple simulated trends.

Our myocardial infarction and kidney disease use-cases show that it is possible to identify well-known concepts for various cell types. This shows that the framework and the approach of analyzing data with the PICASO network is well applicable to real-world data and questions. It is possible to draw connections on originally unconnected concepts, due to the integrated knowledge encoded in the various databases used to construct the PICASO network.

Another limitation of PICASO is that when importing the individual single resources, we had to find a common nomenclature. Focusing on single cell RNA-seq data and a platform agnostic framework, we finally chose to use gene symbols as denominators for nodes representing genes, transcripts and proteins. Most importantly, no distinction between genes, their transcript isoforms and their products, proteins, is made due to lack of available single-cell isoform-specific transcriptomics data.

As the PICASO framework is modular and extensible, it is possible to incorporate more distinction between gene- and protein-expression. Moreover, our framework provides the users flexibility to explore new methods for scoring interactions and nodes. If downstream methods are affected, e.g. because one transitions from scalar scores to vectorized scores, the users can also adapt existing classes for downstream analyses where applicable, e.g. by providing custom score-access functions. Currently, the PICASO framework offers methods which operate on a score based expression system, where gene-expression is converted into aggregate measures such as expression scores, which are the product of mean expression and percent expressed. This can be extended to other distribution-based representations.

Finally, the PICASO framework is a new way of analyzing multi-dimensional single-cell expression datasets. We aggregate existing biomedical knowledge in our PICASO network and provide methods for identifying and interpreting network communities. For characterizing the identified communities, we provide functions and methods in Python as well as provide functionality to use AI-powered functional description using pre-trained LLMs. Overall, PICASO can be used for interpretable hypotheses generation and improving mechanistic understanding of single-cell transcriptomics biomedical data sets.

## CODE AND DATA AVAILABILITY

The source code and notebooks for the analyses presented are available from GitHub https://github.com/mjoppich/PICASO. Processed input files for the PICASO framework will be available from Zenodo upon peer-review.

## AUTHOR CONTRIBUTIONS

MJ and SH conceived the idea and designed the study. MJ carried out the computational analyses and wrote the code. MJ, RK and SH interpreted results and data. MJ and SH wrote the manuscript. All authors contributed to editing the manuscript.

## Supplementary Figures

**Figure S1:**
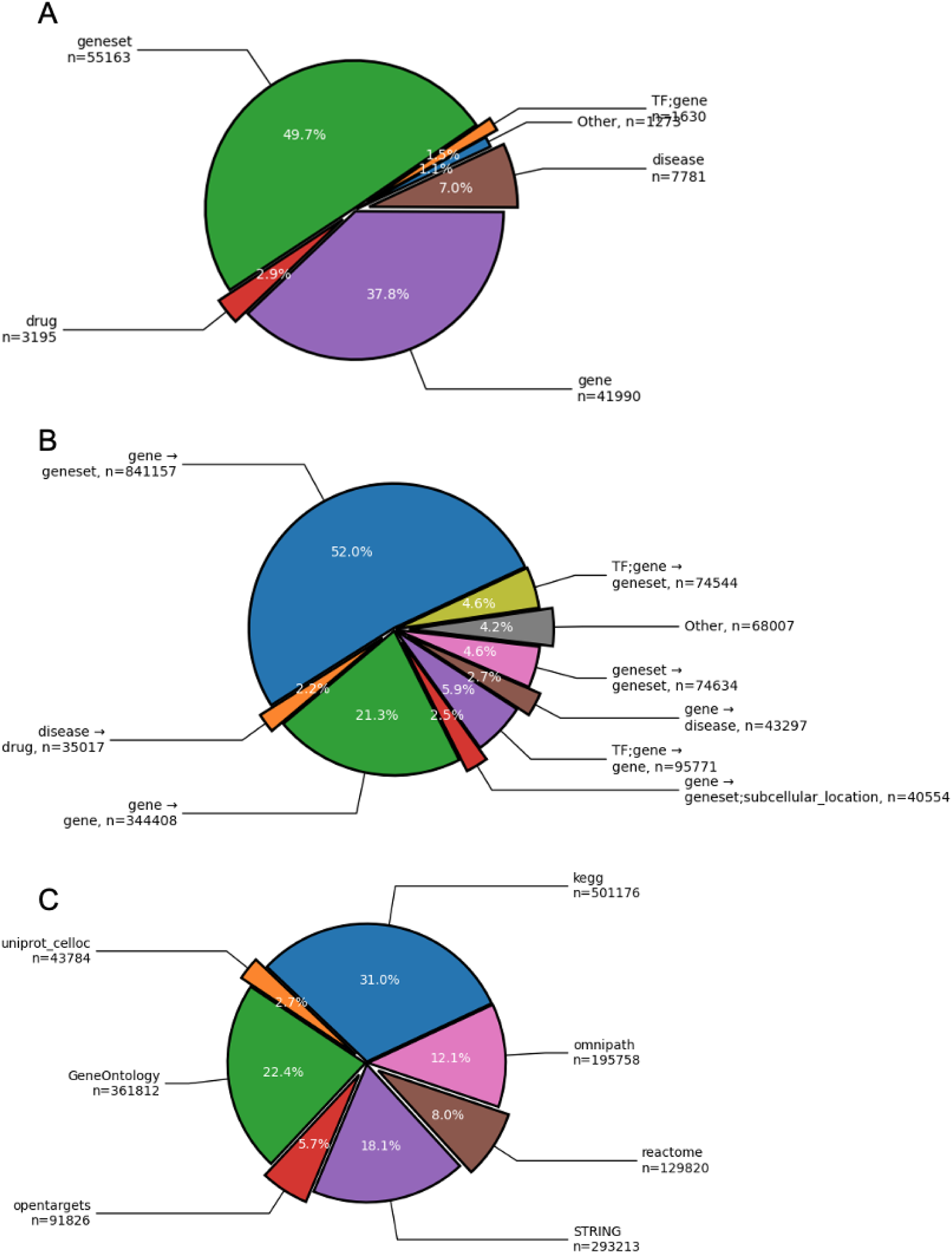
Characterization of underlying public data sets in PICASO. The full PICASO network consists of (A) 111032 many nodes, of which most are of geneset type, followed by gene and disease types. (B) Most of the 1617389 edges are between genes and gene-sets, followed by gene-gene interactions. (C) Various resources are integrated in the PICASO network. Most edges originate from KEGG genesets or Gene Ontology (GO), followed by edges taken from STRING.

**Figure S2:**
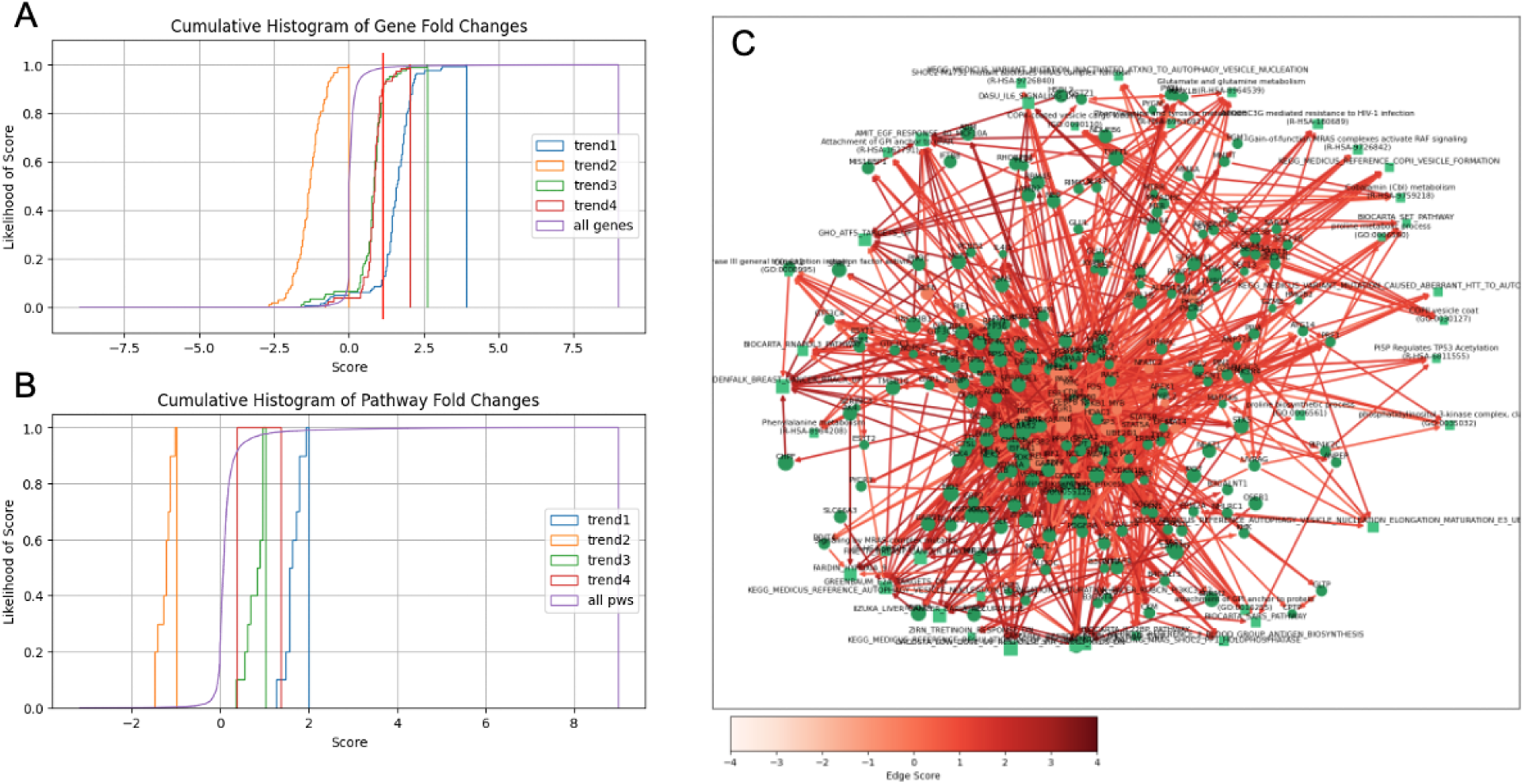
Evaluating PICASO performance on simulated data. (A) Using the differential knowledge graph approach it is possible to identify the relevant genes and (B) pathways. The red line in the gene fold changes shows the fold change of the last ranked gene for further analysis (300th gene). (C) The resulting DNET instance for the simulated trend1 and the comparison of stage1 vs. wild-type.

**Figure S3:**
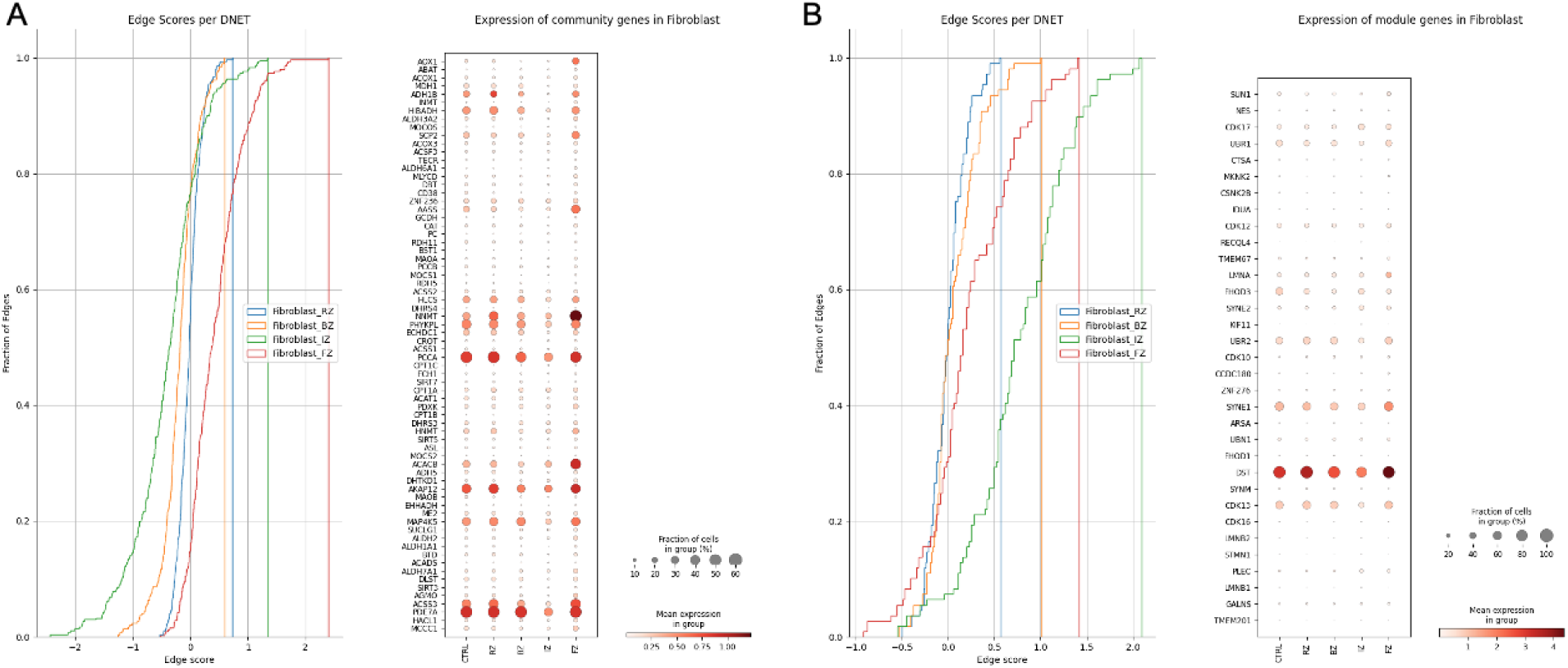
Case-study Myocardial Infarction. A) Module 47, fibroblasts in FZ vs controls - The edge scores per DNET shown as cumulative histogram of the edge scores for all zone-specific DNETs (left). Panel showing the gene expression values in the underlying scRNA-seq data (right). B) Module 127, fibroblasts in IZ vs controls - The edge scores per DNET shown as cumulative histogram of the edge scores for all zone-specific DNETs (left). Panel showing the gene expression values in the underlying scRNA-seq data (right).

**Figure S4:**
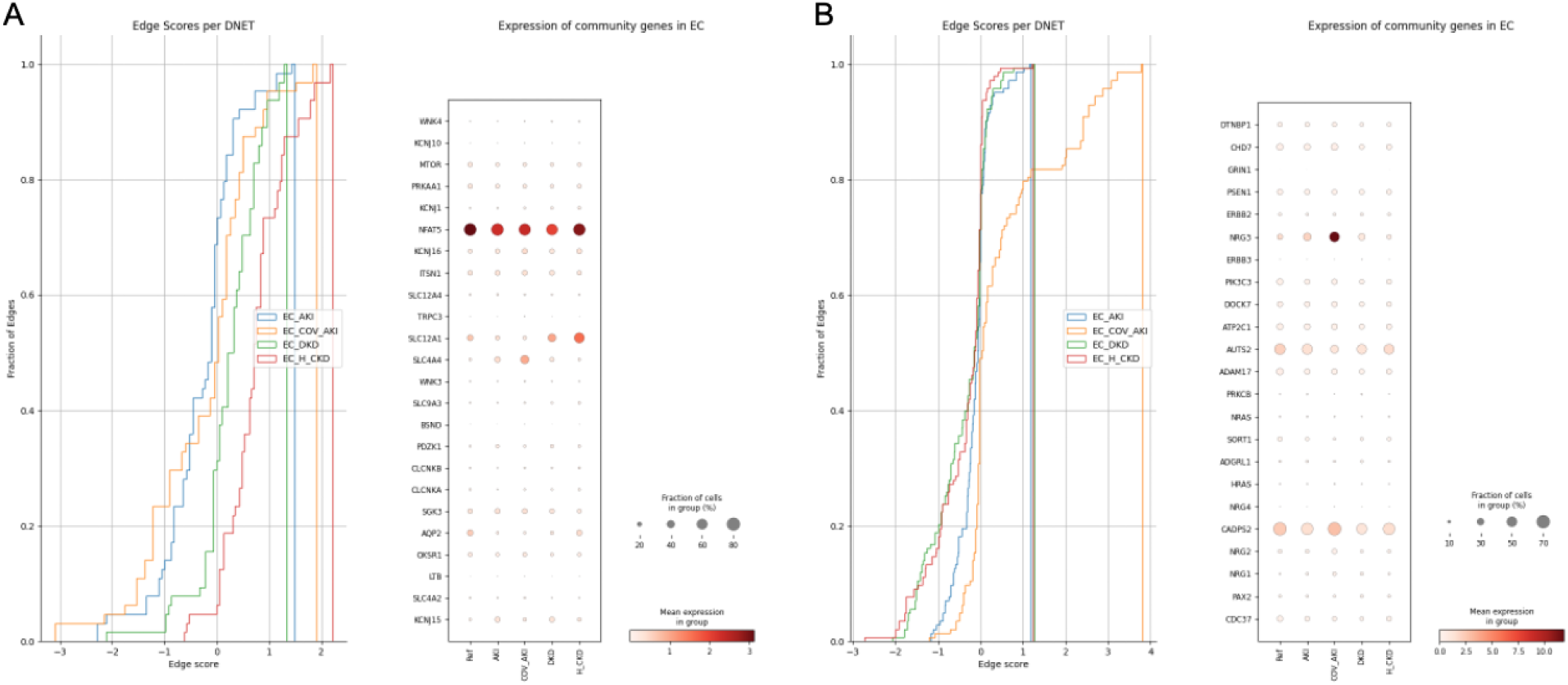
Case-study Kidney Precision Medicine Project. A) Module 14, ECs in H_CKD vs controls - The edge scores per DNET shown as cumulative histogram of the edge scores for all zone-specific DNETs (left). Panel showing the gene expression values in the underlying scRNA-seq data (right). B) Module 29, ECs in COV_AKI vs controls - The edge scores per DNET shown as cumulative histogram of the edge scores for all zone-specific DNETs (left). Panel showing the gene expression values in the underlying scRNA-seq data (right).

## Footnotes

1 https://github.com/slowkow/tftargets

2 https://github.com/cellgeni/sceasy

## REFERENCES

1. Sikkema L, Ramírez-Suástegui C, Strobl DC, Gillett TE, Zappia L, Madissoon E, et al. An integrated cell atlas of the lung in health and disease. Nat Med. 2023;29: 1563–1577.

2. Kuppe C, Ramirez Flores RO, Li Z, Hayat S, Levinson RT, Liao X, et al. Spatial multi-omic map of human myocardial infarction. Nature. 2022;608: 766–777.

3. Kuppe C, Ibrahim MM, Kranz J, Zhang X, Ziegler S, Perales-Patón J, et al. Decoding myofibroblast origins in human kidney fibrosis. Nature. 2021;589: 281–286.

4. Mathys H, Boix CA, Akay LA, Xia Z, Davila-Velderrain J, Ng AP, et al. Single-cell multiregion dissection of Alzheimer’s disease. Nature. 2024. doi:10.1038/s41586-024-07606-7

5. Jiang J, Wang C, Qi R, Fu H, Ma Q. scREAD: A Single-Cell RNA-Seq Database for Alzheimer’s Disease. iScience. 2020;23: 101769.

6. Lake BB, Menon R, Winfree S, Hu Q, Melo Ferreira R, Kalhor K, et al. An atlas of healthy and injured cell states and niches in the human kidney. Nature. 2023;619: 585–594.

7. Abedini A, Levinsohn J, Klötzer KA, Dumoulin B, Ma Z, Frederick J, et al. Single-cell multi-omic and spatial profiling of human kidneys implicates the fibrotic microenvironment in kidney disease progression. Nat Genet. 2024. doi:10.1038/s41588-024-01802-x

8. Pekayvaz K, Losert C, Knottenberg V, Gold C, van Blokland IV, Oelen R, et al. Multiomic analyses uncover immunological signatures in acute and chronic coronary syndromes. Nat Med. 2024;30: 1696–1710.

9. Argelaguet R, Velten B, Arnol D, Dietrich S, Zenz T, Marioni JC, et al. Multi-Omics Factor Analysis-a framework for unsupervised integration of multi-omics data sets. Mol Syst Biol. 2018;14: e8124.

10. Van de Sande B, Lee JS, Mutasa-Gottgens E, Naughton B, Bacon W, Manning J, et al. Applications of single-cell RNA sequencing in drug discovery and development. Nat Rev Drug Discov. 2023;22: 496–520.

11. Lähnemann D, Köster J, Szczurek E, McCarthy DJ, Hicks SC, Robinson MD, et al. Eleven grand challenges in single-cell data science. Genome Biol. 2020;21: 31.

12. Ochoa D, Hercules A, Carmona M, Suveges D, Baker J, Malangone C, et al. The next-generation Open Targets Platform: reimagined, redesigned, rebuilt. Nucleic Acids Res. 2023;51: D1353–D1359.

13. Szklarczyk D, Kirsch R, Koutrouli M, Nastou K, Mehryary F, Hachilif R, et al. The STRING database in 2023: protein–protein association networks and functional enrichment analyses for any sequenced genome of interest. Nucleic Acids Res. 2022;51: D638–D646.

14. Liberzon A, Subramanian A, Pinchback R, Thorvaldsdóttir H, Tamayo P, Mesirov JP. Molecular signatures database (MSigDB) 3.0. Bioinformatics. 2011;27: 1739–1740.

15. Fernández-Torras A, Duran-Frigola M, Bertoni M, Locatelli M, Aloy P. Integrating and formatting biomedical data as pre-calculated knowledge graph embeddings in the Bioteque. Nat Commun. 2022;13: 5304.

16. Lobentanzer S, Aloy P, Baumbach J, Bohar B, Charoentong P, Danhauser K, et al. Democratising Knowledge Representation with BioCypher. arXiv [q-bio.MN]. 2022. Available: http://arxiv.org/abs/2212.13543

17. Chandak P, Huang K, Zitnik M. Building a knowledge graph to enable precision medicine. Sci Data. 2023;10: 67.

18. Sadegh S, Skelton J, Anastasi E, Bernett J, Blumenthal DB, Galindez G, et al. Network medicine for disease module identification and drug repurposing with the NeDRex platform. Nat Commun. 2021;12: 6848.

19. Türei D, Korcsmáros T, Saez-Rodriguez J. OmniPath: guidelines and gateway for literature-curated signaling pathway resources. Nat Methods. 2016;13: 966–967.

20. Kozomara A, Birgaoanu M, Griffiths-Jones S. miRBase: from microRNA sequences to function. Nucleic Acids Res. 2019;47: D155–D162.

21. The Gene Ontology Consortium. Expansion of the Gene Ontology knowledgebase and resources. Nucleic Acids Res. 2017;45: D331–D338.

22. Kanehisa M, Furumichi M, Tanabe M, Sato Y, Morishima K. KEGG: new perspectives on genomes, pathways, diseases and drugs. Nucleic Acids Res. 2017;45: D353–D361.

23. Gillespie M, Jassal B, Stephan R, Milacic M, Rothfels K, Senff-Ribeiro A, et al. The reactome pathway knowledgebase 2022. Nucleic Acids Res. 2022;50: D687–D692.

24. UniProt Consortium. UniProt: a worldwide hub of protein knowledge. Nucleic Acids Res. 2019;47: D506–D515.

25. Hagberg A, Schult D, Swart P, Hagberg JM. Exploring network structure, dynamics, and function using NetworkX. 2008. Available: https://www.osti.gov/biblio/960616

26. García-Lora A, Pedrinaci S, Garrido F. Protein-bound polysaccharide K and interleukin-2 regulate different nuclear transcription factors in the NKL human natural killer cell line. Cancer Immunol Immunother. 2001;50: 191–198.

27. Liberzon A, Birger C, Thorvaldsdóttir H, Ghandi M, Mesirov JP, Tamayo P. The Molecular Signatures Database (MSigDB) hallmark gene set collection. Cell Syst. 2015;1: 417–425.

28. Türei D, Valdeolivas A, Gul L, Palacio-Escat N, Klein M, Ivanova O, et al. Integrated intra-and intercellular signaling knowledge for multicellular omics analysis. Mol Syst Biol. 2021;17: e9923.

29. Wu L, Fan J, Belasco JG. MicroRNAs direct rapid deadenylation of mRNA. Proc Natl Acad Sci U S A. 2006;103: 4034–4039.

30. Eichhorn SW, Guo H, McGeary SE, Rodriguez-Mias RA, Shin C, Baek D, et al. mRNA destabilization is the dominant effect of mammalian microRNAs by the time substantial repression ensues. Mol Cell. 2014;56: 104–115.

31. Wolf FA, Angerer P, Theis FJ. SCANPY: large-scale single-cell gene expression data analysis. Genome Biol. 2018;19: 15.

32. Virshup I, Rybakov S, Theis FJ, Angerer P, Wolf FA. anndata: Annotated data. BioRxiv. 2021. Available: https://www.biorxiv.org/content/10.1101/2021.12.16.473007.abstract

33. Piñero J, Ramírez-Anguita JM, Saüch-Pitarch J, Ronzano F, Centeno E, Sanz F, et al. The DisGeNET knowledge platform for disease genomics: 2019 update. Nucleic Acids Res. 2020;48: D845–D855.

34. Virtanen P, Gommers R, Oliphant T, Haberland M, Reddy T, Cournapeau D, et al. SciPy 1.0: fundamental algorithms for scientific computing in Python. Nat Methods. 2019;17: 261–272.

35. Ahn Y-Y, Bagrow JP, Lehmann S. Link communities reveal multiscale complexity in networks. Nature. 2010;466: 761–764.

36. Van Dongen S. Graph Clustering Via a Discrete Uncoupling Process. SIAM J Matrix Anal Appl. 2008;30: 121–141.

37. Kheirkhahzadeh M, Lancichinetti A, Rosvall M. Efficient community detection of network flows for varying Markov times and bipartite networks. Phys Rev E. 2016;93: 032309.

38. Smiljanić J, Blöcker C, Holmgren A, Edler D, Neuman M, Rosvall M. Community Detection with the Map Equation and Infomap: Theory and Applications. arXiv [physics.soc-ph]. 2023. Available: http://arxiv.org/abs/2311.04036

39. Peng J, Uygun S, Kim T, Wang Y, Rhee SY, Chen J. Measuring semantic similarities by combining gene ontology annotations and gene co-function networks. BMC Bioinformatics. 2015;16: 44.

40. huggingface_hub: The official Python client for the Huggingface Hub. Github; Available: https://github.com/huggingface/huggingface_hub

41. Andrei. llama-cpp-python: Python bindings for llama.cpp. Github; Available: https://github.com/abetlen/llama-cpp-python

42. Creators Mueller AC. Wordcloud. doi:10.5281/zenodo.10321882

43. Baruzzo G, Patuzzi I, Di Camillo B. SPARSim single cell: a count data simulator for scRNA-seq data. Bioinformatics. 2020;36: 1468–1475.

44. Han H, Cho J-W, Lee S, Yun A, Kim H, Bae D, et al. TRRUST v2: an expanded reference database of human and mouse transcriptional regulatory interactions. Nucleic Acids Res. 2018;46: D380–D386.

45. Cellxgene Data Portal. In: Cellxgene Data Portal [Internet]. [cited 19 Jul 2024]. Available: https://cellxgene.cziscience.com/collections/bcb61471-2a44-4d00-a0af-ff085512674c

46. Padda IS, Mahtani AU, Parmar M. Sodium-Glucose Transport Protein 2 (SGLT2) Inhibitors. StatPearls Publishing; 2023.

47. Kaludercic N, Mialet-Perez J, Paolocci N, Parini A, Di Lisa F. Monoamine oxidases as sources of oxidants in the heart. J Mol Cell Cardiol. 2014;73: 34–42.

48. Deluyker D, Ferferieva V, Driesen RB, Verboven M, Lambrichts I, Bito V. Pyridoxamine improves survival and limits cardiac dysfunction after MI. Sci Rep. 2017;7: 16010.

49. Ahadome SD, Abraham DJ, Rayapureddi S, Saw VP, Saban DR, Calder VL, et al. Aldehyde dehydrogenase inhibition blocks mucosal fibrosis in human and mouse ocular scarring. JCI Insight. 2016;1: e87001.

50. Chen Y, Du J, Zheng L, Wang Z, Zhang Z, Wu Z, et al. Chemical screening links disulfiram with cardiac protection after ischemic injury. Cell Regen. 2023;12: 25.

51. Yang H, Shi Y, Liu H, Lin F, Qiu B, Feng Q, et al. Pyroptosis executor gasdermin D plays a key role in scleroderma and bleomycin-induced skin fibrosis. Cell Death Discov. 2022;8: 183.

52. Killalea SM, Krum H. Systematic review of the efficacy and safety of perhexiline in the treatment of ischemic heart disease. Am J Cardiovasc Drugs. 2001;1: 193–204.

53. Labrak Y, Bazoge A, Morin E, Gourraud P-A, Rouvier M, Dufour R. BioMistral: A Collection of Open-Source Pretrained Large Language Models for Medical Domains. arXiv [cs.CL]. 2024. Available: http://arxiv.org/abs/2402.10373

54. LoneStriker/BioMistral-7B-DARE-GGUF · Hugging Face. [cited 30 Jul 2024]. Available: https://huggingface.co/LoneStriker/BioMistral-7B-DARE-GGUF

55. LifeMap Sciences. Emery-Dreifuss Muscular Dystrophy - MalaCards. [cited 20 Jul 2024]. Available: https://www.malacards.org/card/emery_dreifuss_muscular_dystrophy

56. Hsu A, Duan Q, Day DS, Luo X, McMahon S, Huang Y, et al. Targeting transcription in heart failure via CDK7/12/13 inhibition. Nat Commun. 2022;13: 4345.

57. Rath O, Kozielski F. Kinesins and cancer. Nat Rev Cancer. 2012;12: 527–539.

58. Bahleda R, Grilley-Olson JE, Govindan R, Barlesi F, Greillier L, Perol M, et al. Phase I dose-escalation studies of roniciclib, a pan-cyclin-dependent kinase inhibitor, in advanced malignancies. Br J Cancer. 2017;116: 1505–1512.

59. Thomas TP, Grisanti LA. The Dynamic Interplay Between Cardiac Inflammation and Fibrosis. Front Physiol. 2020;11: 529075.

60. Mercer PF, Chambers RC. Coagulation and coagulation signalling in fibrosis. Biochim Biophys Acta. 2013;1832: 1018–1027.

61. Konrad M, Nijenhuis T, Ariceta G, Bertholet-Thomas A, Calo LA, Capasso G, et al. Diagnosis and management of Bartter syndrome: executive summary of the consensus and recommendations from the European Rare Kidney Disease Reference Network Working Group for Tubular Disorders. Kidney Int. 2021;99: 324–335.

62. Starremans PGJF, Kersten FFJ, Knoers NVAM, van den Heuvel LPWJ, Bindels RJM. Mutations in the human Na-K-2Cl cotransporter (NKCC2) identified in Bartter syndrome type I consistently result in nonfunctional transporters. J Am Soc Nephrol. 2003;14: 1419–1426.

63. Welling PA, Ho K. A comprehensive guide to the ROMK potassium channel: form and function in health and disease. Am J Physiol Renal Physiol. 2009;297: F849–63.

64. Keck M, Andrini O, Lahuna O, Burgos J, Cid LP, Sepúlveda FV, et al. Novel CLCNKB mutations causing Bartter syndrome affect channel surface expression. Hum Mutat. 2013;34: 1269–1278.

65. St Clair EW, Baer AN, Wei C, Noaiseh G, Parke A, Coca A, et al. Clinical Efficacy and Safety of Baminercept, a Lymphotoxin β Receptor Fusion Protein, in Primary Sjögren’s Syndrome: Results From a Phase II Randomized, Double-Blind, Placebo-Controlled Trial. Arthritis Rheumatol. 2018;70: 1470–1480.

66. O’shea J, Tato CM, Siegel R. Cytokines and cytokine receptors. Clinical Immunology. Elsevier; 2008. pp. 139–171.

67. Moreno J, Gluud LL, Galsgaard ED, Hvid H, Mazzoni G, Das V. Identification of ligand and receptor interactions in CKD and MASH through the integration of single cell and spatial transcriptomics. PLoS One. 2024;19: e0302853.

68. Dugaucquier L, Feyen E, Mateiu L, Bruyns TAM, De Keulenaer GW, Segers VFM. The role of endothelial autocrine NRG1/ERBB4 signaling in cardiac remodeling. Am J Physiol Heart Circ Physiol. 2020;319: H443–H455.

69. Saul S, Karim M, Ghita L, Huang P-T, Chiu W, Durán V, et al. Anticancer pan-ErbB inhibitors reduce inflammation and tissue injury and exert broad-spectrum antiviral effects. J Clin Invest. 2023;133. doi:10.1172/JCI169510

70. Oubounyt M, Elkjaer ML, Laske T, Grønning AGB, Moeller MJ, Baumbach J. De-novo reconstruction and identification of transcriptional gene regulatory network modules differentiating single-cell clusters. NAR Genom Bioinform. 2023;5: lqad018.

71. Bravo González-Blas C, De Winter S, Hulselmans G, Hecker N, Matetovici I, Christiaens V, et al. SCENIC+: single-cell multiomic inference of enhancers and gene regulatory networks. Nat Methods. 2023;20: 1355–1367.

72. Geistlinger L, Csaba G, Küffner R, Mulder N, Zimmer R. From sets to graphs: towards a realistic enrichment analysis of transcriptomic systems. Bioinformatics. 2011;27: i366–73.

73. Forelli N, Eaton D, Patel J, Bowman CE, Kawakami R, Kuznetsov IA, et al. SGLT2 inhibitors activate pantothenate kinase in the human heart. bioRxiv. 2024. doi:10.1101/2024.07.26.605401

